# Interactions Underlying Stress Granule Structure and Therapeutic Dissolution

**DOI:** 10.1101/2025.03.10.642463

**Authors:** Jay L. Kaplan, Michael A. Webb

## Abstract

Stress granules are biomolecular condensates composed of RNA and proteins that form in response to stress; their dysregulation is implicated in neurodegenerative diseases. In this study, we develop a minimal stress-granule model, composed of RNA and six key proteins associated with neurodegenerative conditions, and study its characteristics using coarse-grained molecular dynamics simulations. We find that RNA is essential to form stable condensates in these biopolymer mixtures, while underlying protein-protein interactions result in heterogeneous, multi-phasic architectures. Inspired by therapeutic applications, we then challenge the stability of these condensates in the presence of twenty distinct small molecules. Simulation-derived properties classify compounds as “dissolving” or “non-dissolving” with 85% agreement with experimental findings. Further analysis suggests that dissolving compounds disrupt stress granule structure by preferentially associating with RNA and stripping the scaffold that maintains its multiphasic architecture. These insights advance understanding of stress granule stability and demonstrate modeling strategies for screening of therapeutic candidates.

## INTRODUCTION

Stress Granules (SGs) are biomolecular condensates, membrane-less organelles that form through liquid-liquid phase separation, featuring ribonucleoprotein assemblies characterized by a vast proteomic and transcriptomic network^1,2^. SGs form in the cytoplasm upon exposure to environmental stressors and possess biphasic morphologies with dense cores surrounded by a more diffuse shell^3^. SG formation is thought to be driven by certain protein species that interact with mRNA and sequester it inside a condensate, thereby controlling mRNA metabolism and translation during cellular stress^4,5^ by either storing mRNA, transferring it for degradation, or releasing it for polysomes to re-initiate translation^6^. Chronic cellular stress, mutations in key proteins, alterations in various mRNA transcripts, and sufficiently high concentrations of specific proteins prevent dissolution of SGs and can cause the formation of aberrant, persistent SGs, which promote further aggregation of disease-related proteins. These aberrant SGs and the accompanying aggregation of disease-related proteins and mRNA transcripts have been linked to various diseases, including neurodegenerative disorders, cancers, and viral infections, making these aberrant condensates a promising target for novel therapeutics^7^.

There is substantial interest in small-molecule compounds as prospective therapeutics that function by modulating the phase behavior and stability of biomolecular condensates, including SGs^8–10^. SG formation is thought to be driven by the macromolecular crowding of accumulated mRNA and proteins. This enables multivalent interactions between the mRNA and key client and scaffold proteins^11^. The excessive aggregation of specific proteins within SGs, such as TDP43 and FUS, can cause the transition of the condensate from a liquid-like to solid-like state. This inhibits disassembly and leads to subsequent chronic SG formation with consequent further aggregation of these disease-inducing biopolymers in the affected cells^12,13^. Therefore, understanding SG structure, dynamics, and stability could reveal many therapeutic targets^14^. A limited number of experimental studies have screened existing small-molecule libraries to determine which compounds might partition into, and modulate, SG stability^10,15,16^. Empirical observations suggest that particular physicochemical factors and mechanisms of interaction may be leveraged to promote SG dissolution. For example, many small-molecule compounds with notable “dissolving” effects on SGs empirically possess aromatic side chains^16^. Additionally, other molecule-small compounds apparently prevent TDP43 localization and accumulation into SGs as well as the formation of cytoplasmic puncta^16^, a hallmark of amyotrophic lateral sclerosis (ALS)^17^. Although there may be well-reasoned hypotheses regarding how and why certain small-molecule compounds effectively interact with SGs, direct evidence via experimental characterization can be challenging to obtain in such complex systems^18^. Developing principled and alternative methods to rapidly investigate the interactions of small-molecule compounds with SGs would bolster efforts in condensate-modulating therapeutic development.

Molecular dynamics (MD) simulations can offer molecular-level insights into complex systems and phenomena, including phase-separating disordered proteins^19,20^. Due to the range of spatiotemporal scales in the underlying biophysics, multiscale simulation and coarse-grained (CG) modeling^21,22^ have been particularly useful for providing insights into both single-chain and condensed-phase systems composed of intrinsically disordered proteins and RNA^23–33^. However, for residue-level or higher-resolution models, simulation of multicomponent biomolecular condensates, including stress-granule-like systems, is limited but could offer valuable insights into their properties^34,35^. By extension, simulation studies on the interactions and effects of small-molecule compounds on biomolecular condensate properties are nascent. Therefore, the efficacy of using CG modeling to study SG structure and dissolution remains unexplored but offers significant applications in structural biology and high-throughput *in silico* screens for biomolecular condensate modulating therapeutics^8^.

Here, we study the structure and stability of SGs, including their interactions with small-molecule compounds, using data-driven CG modeling. We first generate a computationally tractable model with biophysical relevance to stress granules and examine its structural properties and underlying interactions. Using a simple and efficient data-driven coarse-graining parameterization procedure, we then investigate how properties of SGs are impacted by a chemically diverse set of small-molecule compounds (SMCs). Our simulation results agree with experimental outcomes for 85% of the SMCs regarding their propensity to modulate SG stability. Despite the small system size relative to *in vivo* SGs and the coarse-grained resolution afforded by the MPiPi model used in these simulations, we elucidate additional aspects of SG structure and identify possible mechanisms of SMC-induced stress-granule destabilization. This includes the preferential association of SMCs with RNA that reduces protein-RNA contacts, which induces intra-condensate mixing and a collapse of the SG’s multiphasic architecture. Overall, this work provides new molecular-level understanding of stress-granule properties as well as a platform for rapidly screening and developing biomolecular condensate-modulating SMC candidates.

## RESULTS AND DISCUSSION

### Minimal mixtures of SG proteins and RNA yield stable condensates

Stress granules are highly complex non-membrane-bound organelles with a diverse proteome consisting of hundreds of associated proteins^3,36^. Therefore, we first aim to identify a computationally tractable model for the core of a SG, which is composed of both proteins and mRNA.

For the proteins, we consider a reduced proteome composed of only Tier 1 SG proteins, defined as protein species that possess high affinity to localize into SGs^**?**^ . From this subset, we select proteins strongly implicated in neurodegenerative disorders^37^ and then isolate only those proteins that have 800 or fewer amino acids (to reduce computational complexity). This results in a set of six proteins: G3BP1, FUS, TIA1, TTP, TDP43, and PABP1. Figure 1A shows the distribution of amino acids in these proteins, indicating a diverse range of possible interactions. Based on the relatively higher concentration of G3BP1 found in SG cores and its key role in SG formation^11,38,39^, we set the number of G3BP1 proteins to be approximately twice that of all other proteins. For simplicity and to preserve RNA Recognition Motifs (RRMs), we use the full-length protein sequences, including both ordered and disordered regions. Representing ordered regions as disordered with a force field developed for IDPs may enhance the valency of the biopolymers and thereby the tendency to phase separate, though excluding the structured domains entirely would introduce different approximations. Because the aim of this work is to compare how small-molecule compounds perturb the condensate, the relevant quantities are differences taken within a consistent modeling framework against a deliberately stable reference SG, for which such a systematic effect is largely mitigated. Accurately modeling both ordered and disordered regions is an active area of research,^27,40–44^ and future studies may examine its implications for the devised computational SG.

**Figure 1.**
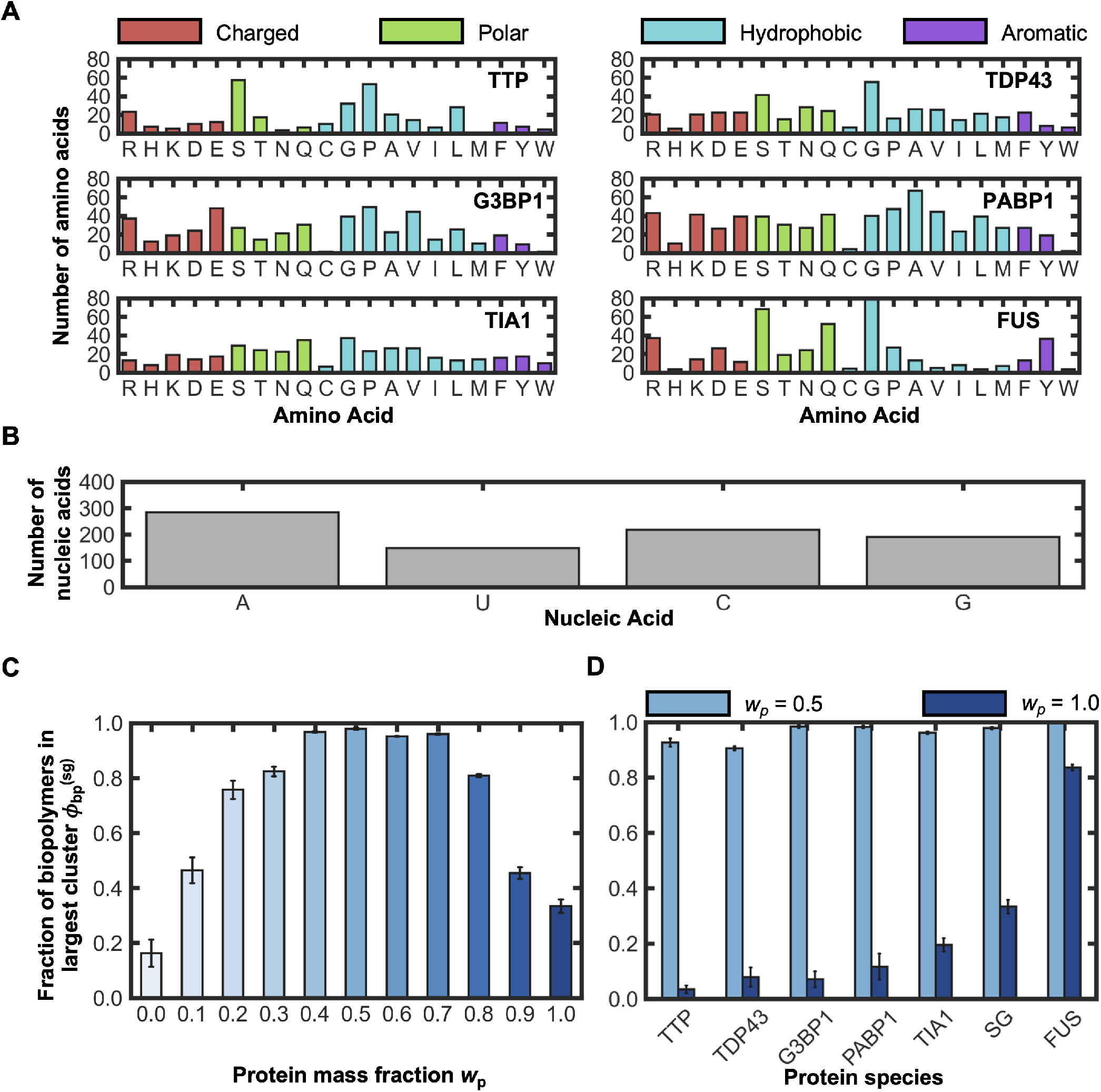
A mixture of key proteins and RNA form stable condensates, providing a simple model for the core of a stress-granule. **(A)** Distributions of amino acids in proteins present in the model stress granule: TTP, G3BP1, TIA1, TDP43, PABP1, and FUS. The amino acids are color-coded based on their functional group classification. **(B)** Distribution of nucleic acids in the modified ZACN poly-(A) mRNA transcript present in the model stress granule. **(C)** Examination of the fraction of biopolymers within the largest cluster, 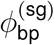, identified in systems with different mass fractions of protein, *w*_p_. The color of the bars also indicates the mass fraction of the protein present in the system with darker bars representing a larger *w*_p_ . **(D)** Comparison of the fraction of biopolymers within the largest cluster, 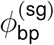, for different minimal SG models of a single protein species for systems with the optimal mass fraction of (*w*_p_ = 0.5) and without (*w*_p_ = 1.0) RNA.

For the mRNA, we form a single, relatively short RNA transcript. In particular, we begin with the ZACN transcript, which exhibits a high affinity to localize into SGs^2^. We then extract only a randomized portion of the coding region and exclude the UTR regions to reduce the transcript length from 2,651 to 700 nucleic acids. We then add a 140-nucleotide poly-(A) tail, as SGs stain positive for poly-(A) mRNA^2^. Figure 1B shows the distribution of nucleic acids for the model RNA sequence. The simplifications described may be reasonable because the diverse SG transcriptome does not require specific mRNA transcripts or characteristics to phase separate^45^. However, RNA strand length and sequence can still influence the phase-separating behavior of RNP granules^2,46,47^, which motivates future consideration of how RNA composition and length impact the observations from computational SG models.

To determine whether mixtures of the aforementioned biopolymers can form stable condensates, we perform 2 µs MD simulations with a 50 ns equilibration period using the MPiPi force field^27^. This force field utilizes an implicit solvent and treats amino and nucleic acids at a one-bead-per-residue resolution; interactions are modeled via bonded and non-bonded terms, including explicit electrostatics with screening (see also “Details of MD Simulations”). It is important to note that this model does not accurately represent complex structured protein regions but effectively represents IDPs and LLPS while allowing for running simulations of sufficient scale to investigate SG properties. Effectively modeling both structured domains and intrinsically disordered regions (IDRs) is an existing challenge for coarse-grained protein force fields^48^, and given the strong presence of IDPs in SGs, we elected a model effective for the latter. Therefore, the results focus on the general propensity to phase separate based on the proportion and sequence of amino acids and on proteins with long Intrinsically Disordered Regions (IDRs).

We quantify stability of a biopolymer mixture by the fraction of biopolymers found in the largest connected aggregate in the system, 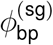, and the total number of individual aggregates, *k*; the expectation is that more stable SGs will possess larger 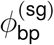 and smaller *k*. Figure 1C shows that stable condensates form over a range of intermediate protein mass fractions, *w*_p_ (defined as the mass of protein in the system relative to the total mass of biopolymer). In pure RNA systems (*w*_p_ = 0.0), phase-separation of the RNA is limited, with 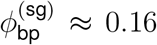. The negligible 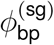 is due to transient interactions among RNA strands; RNA can also phase-separate in the absence of proteins under certain conditions^49^. However, once the system contains a nominal protein fraction (e.g., *w*_p_ = 0.1), 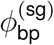 rises to ca. 0.48, approximately three-fold over the pure RNA systems. This indicates greater propensity for a protein-RNA mixture to induce liquid-liquid phase separation and that RNA is a key scaffold of SGs. Nearly all biopolymers interact to form a single large cluster for protein mass fractions of 0.4 ≥ *w*_p_ ≥ 0.7. These values align with prior work using phenomenological models^50^, though some experiments suggest greater mRNA composition^2^. The disparity is possibly related to the use of our comparatively short RNA transcript which reduces the number of available sites for RNA-protein interactions and motivates further investigation on the effect of RNA length and composition on SG stability. Nevertheless, we characterize the system with *w*_p_ = 0.5 as a “control” SG model, as it represents the most stable condensate and therefore a challenging test for dissolution by SMCs.

### Heterotypic protein-RNA interactions underlie SG formation

To better understand drivers of stability in the SG model, we contrast systems consisting of each individual composite protein species with those consisting of the full SG mixture both in the presence of RNA, specifically the optimal protein mass fraction, *w*_p_ = 0.5, and in the absence of RNA, *w*_p_ = 1.0. Figure 1D shows that RNA is essential to form the model SG, with 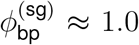 for all protein species systems containing the optimal RNA mass fraction. This is compared to systems in the absence of RNA that exhibit 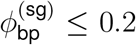 for five of the six protein systems, 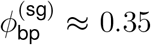 for the SG mixture, and 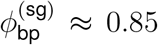 for the pure FUS system. These results broadly agree with prior expectations regarding the role of mRNA in SGs^5,47,51,52^ and also essential components in the SG proteome^3,11,38^. In particular, five of the six protein species do not centralize into a significant single cluster in the absence of RNA (i.e., *w*_p_ = 1.0); the lone exception is FUS, which is known to phase-separate without RNA. That five of the six species fail to condense on their own indicates that representing ordered regions as disordered does not by itself drive spurious assembly, although it may still enhance the interactions of these proteins with RNA. Without RNA, the propensity for protein localization aligns well with the overall abundance of aromatic residues, particularly Tyrosine, which perhaps highlights the relevance of aromatic interaction grammar deemed vital for LLPS^27,53^. With RNA (i.e., *w*_p_ = 0.5), virtually all biopolymers coalesce into a single droplet. Comparing the full SG system to systems of individual composite proteins, the protein mixture (labeled as ‘SG’ in Figure 1D) exhibits the second highest propensity for aggregation without RNA (after FUS). This suggests that heterotypic (mixed species) protein interactions for this minimal SG model are a stronger driver of condensate formation than homotypic (same species) interactions; protein-RNA interactions are specifically impactful.

### SGs exhibit a hierarchically heterogeneous morphology

Having established a stable, minimalist model SG, we now examine its structural characteristics. Figure 2A qualitatively illustrates the heterogeneous distribution of composite biopolymers within a representative configuration of the model SG. The structure possesses a dense protein-RNA mixture at the center surrounded by a more diffuse, RNA-rich shell. Figure 2B offers quantitative confirmation, examining the distribution of biopolymers and overall size of the model SG via radial density profiles (RDPs). The dense protein-RNA core has an overall concentration of 464 mg/mL and a protein concentration of 340 mg/mL, consistent with prior protein condensate literature^54–56^. The core concentration remains relatively constant up to about 100 Å before decreasing, signifying the transition between the concentrated condensate (SG) and dilute bulk phases. The ratio of protein to RNA concentrations inverts around 250 Å (zoomed inset in Figure 2B), indicating the transition to the diffuse RNA-rich shell.

**Figure 2.**
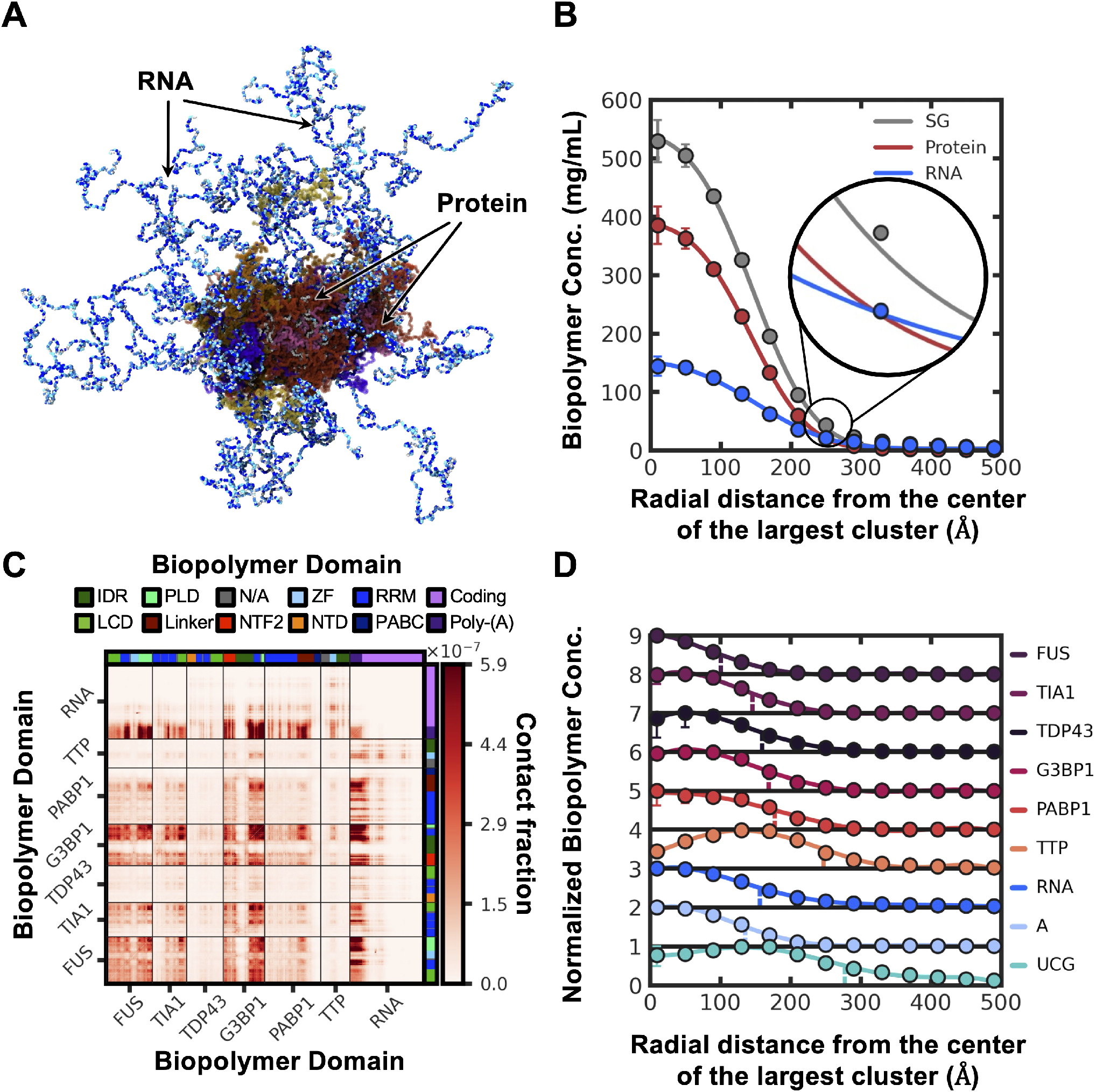
The model stress granule possesses a heterogeneous structure. **(A)** Representative simulation snapshot following equilibration; RNA chains splay outward while proteins concentrate in a dense core. **(B)** Total radial mass-density profile measured from the largest-cluster center of mass, resolved by biopolymer class (whole SG, all protein, all RNA). Inset: magnified view of the condensate edge where the protein and RNA density profiles cross. **(C)** Domain-resolved contact map of the stress granule. Each chain is decomposed into its constituent sequence domains (legend: IDR, PLD, LCD, Linker, NTF2, NTD, ZF, RRM, PABC, and the Poly-(A)/Coding RNA segments), grouped by biopolymer species. Color encodes the *contact fraction*, the share of all inter-chain biopolymer contacts carried by each domain pair (sequential scale, darker = more contacts). **(D)** normalized radial mass-density profiles resolved by biopolymer species, with RNA further split into its poly-(A) (A) and coding (UCG) nucleotides. Each curve is normalized to its own maximum concentration and offset vertically by ‘1’ for clarity.

Figure 2C presents a domain-resolved contact map in which each chain is split into its constituent sequence domains, showing that contacts concentrate among the protein domains and the RNA poly-(A) segment that occupy the dense core. Figure 2D resolves the radial density profile by biopolymer species and indicates that different biopolymers preferentially localize to specific SG regions. This is revealed by tracking visual landmarks, such as the distances from the center of the SG at maximum and half-maximum concentrations. For example, FUS has the highest central concentration and the smallest mean radial position, making it the most centrally localized protein and the SG core anchor. TDP43, by contrast, is the sparsest protein and, although its low concentration is weakly peaked near the center, is distributed broadly throughout the condensate; by the mean radial position of its chains it is among the least centrally localized proteins (Supplementary Information, Figure S10). Meanwhile, G3BP1 maintains a relatively constant concentration up to 90 Å, then gradually decreases to half-maximum at 160 Å. This is consistent with observations of TDP43 de-mixing from G3BP1 in stress granules^57^ and of multiphasic architectures in two protein and RNA systems^31^ but has not been observed in a system of this complexity. The other proteins—TIA1, PABP1, and TTP—are lower in concentration and more broadly distributed. TIA1 forms a broad, low plateau near the center, and PABP1 likewise peaks centrally but declines only slowly, so that its mass extends to intermediate radii; TTP is the most dilute and lies farthest out among the proteins, nearest the diffuse RNA boundary. Up to its half-maximum concentration at 170 Å, RNA distribution closely tracks G3BP1 but decays thereafter. This altogether suggests that the model SG has a dense core of heterogeneously distributed proteins mixed with RNA, surrounded by a diffuse, RNA-rich shell, indicating a multi-phasic structure.

Within Figure 2D, the RNA is further resolved into its poly-(A) and coding (UCG) nucleotides, which are also heterogeneously distributed: the normalized poly-(A) signal is maximal near the SG center (around 90 Å) before decreasing sharply, whereas the coding nucleotides are lower than poly-(A) at the center but exceed it by about 180 Å. The data indicate that RNA adopts a preferential orientation, with the poly-(A) tail embedded in the dense protein–RNA core and the coding regions extending into the diffuse shell. This suggests that the RNA exhibits amphiphilic character, interacting with both core and peripheral species. Previous studies have shown that RNA length influences phase behavior, with shorter transcripts acting as surfactants and longer ones as stabilizers^46^. These simulations suggest that the modeled RNA transcript may function as both. Overall, Figure 2 highlights multiple levels of structural organization within the model SG.

### Biopolymer interactions are varied and specific in the SG

To characterize the hierarchical SG structure, we calculated the frequency of biopolymer interactions using contact maps, resolved by species (Figures 3A,B) and by residue type (Figures 3C,D). Both the contact fraction (Figures 3A,C), giving each pair’s share of all inter-chain contacts, and the log contact enrichment (Figures 3B,D), giving the contact frequency relative to a null model, are analyzed; the enrichment maps reveal interaction trends based on affinity rather than abundance. Figure 3 overall illustrates that varied interaction patterns among biopolymers underlie the observed multiphasic structure of the SG.

**Figure 3.**
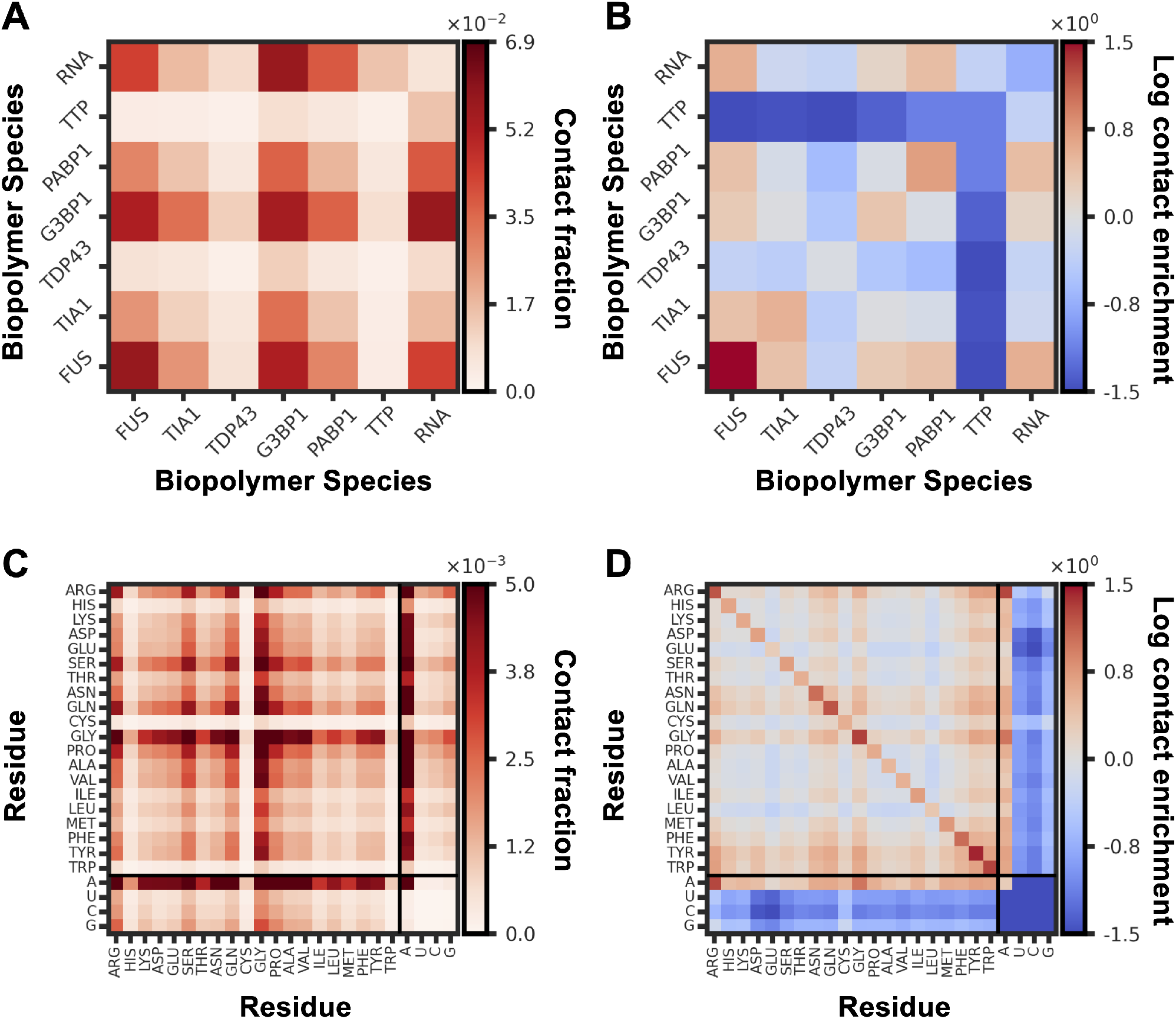
Interaction patterns between protein, RNA, and their amino/nucleic acids underlie the structure of the model stress granule. All maps show inter-chain contacts only. **(A)** Contact fraction between biopolymer species: the share of all inter-chain contacts carried by each species pair (sequential scale, darker = more contacts). **(B)** Log contact enrichment between biopolymer species, ln(*f/q*), where *f* is the observed contact fraction in (A) and *q* the fraction expected under a chain-pair null model (chains paired at random, independent of chain length). Diverging scale centered at 0: red = contacts more frequent than expected, blue = less frequent. **(C)** Contact fraction resolved by amino-acid and nucleic-acid identity. **(D)** Log contact enrichment resolved by amino-acid/nucleic-acid identity, with *q* set by a residue-abundance null model (pairs drawn in proportion to each residue type’s abundance in the condensate). Diverging scale centered at 0.

Examining interactions resolved by species, Figure 3A supports the observation that RNA serves as a scaffold in the SG model. It is essential for the formation of the condensate and exhibits high contact probabilities with most proteins, except for TDP43 and TTP. We hypothesize that disruption of these RNA-protein interactions functions to dissolve SGs. G3BP1, a core SG network protein^11,38,39^, exhibits strong interactions with all biopolymers, especially with itself and RNA. TDP43, which is sparse and de-mixes within the SG, shows low contact probabilities overall but interacts more with G3BP1, RNA, and FUS, while having minimal interaction with TTP, which resides in the SG periphery. The log contact enrichment in Figure 3B highlights that biopolymers have a higher affinity to engage in homotypic interactions but still exhibit significant heterotypic contacts with RNA that are necessary for condensate formation, supporting prior observations of intra-condensate de-mixing^31,57^. FUS shows the strongest homotypic interactions and significant contacts with RNA, consistent with its phase-separation behavior^54,58^. TDP43 also exhibits strong self-interactions and de-mixes into a self-associated sub-region within the SG^57^; the domain-resolved contact map (Figure 2C) shows high contact frequencies among its IDR, RRM, and NTD domains, consistent with previous studies^59^.Because Figure 2C places all species on a single shared contact scale, TDP43’s relatively sparse self-contacts appear muted; resolved on a per-species scale (Supplementary Information, Figure S13), TDP43’s IDR/RRM/NTD self-contact architecture is clearly preserved. Proteins like TIA1 and G3BP1 interact significantly with FUS and RNA, with G3BP1 driving phase-separation via poly-(A) RNA^38,60^. The RNA itself shows homotypic interactions, likely due to poly-(A) tail embedding, while PABP1 and TTP exhibit strong contacts with RNA in line with their known binding behaviors^61^. This reflects an intriguing balance, as the presence of both RNA and protein is critical for SG formation (Figure 1C) and that multiple protein species systems more readily form condensates than single protein species systems (Figure 1D), yet once formed, intra-condensate demixing results in a multiphasic architecture (Figure 2C) arising predominantly from homotypic interactions (Figures 3A,B).

The residue-resolved contact maps in Figures 3C,D suggest that a subset of residues play central roles in determining SG structure. In the contact-fraction maps (Figure 3C), which report each pair’s share of all inter-chain contacts, Arg, Ser, Gln, Gly, and Pro form many contacts, reflecting both abundance and a tendency to interact. When expressed as the log contact enrichment relative to a residue-abundance null model (Figure 3D), Asn, Gln, Gly, Tyr, and Trp display the highest contact propensities. Several of these residues, including Arg, Gln, Tyr, and Trp, have been noted to promote phase separation through hydrogen–bond, *π*–*π*, and electrostatic interactions^53,62–64^. Although such interactions are not directly represented by the CG model, which can only include such effects effectively through the parameterized pair potentials, the contact patterns may be suggestive of their relevance to the SG structure. In contrast, Gly and Pro, which are generally viewed as flexible spacers, may play a supporting structural role. Furthermore, proteins in the SG model, such as FUS, TIA1, PABP1, TDP43, TTP, and G3BP1, are rich in these residues^60,65–68^, reinforcing their roles in phase separation. Strong contacts of Arg and Gly with adenine, prominent due to the poly-(A) tail, align with known RNA-binding motifs^69,70^, and normalized maps highlight the dominance of RNA-RNA C, G and U interactions.

Altogether, analysis of the contact maps supports both roles of RNA: the poly-(A) tail embedded in the core serving as a scaffold and the coding regions in the diffuse shell serving as a surfactant. This is further illustrated in the domain-resolved contact map, which tracks sequence region rather than residue type (Figure 2C). Trp homotypic interactions, driven by *π*-*π* bonding, remain significant despite the low abundance of Trp. These findings, combined with adenine core localization, reflect the expected biophysical behavior of SGs, confirming the relevance of the model to previous experimental insights.

### Simulations effectively classify dissolving and non-dissolving compounds

We now evaluate the impact of small-molecule compounds (SMCs) using MD simulations to measure the effect of each CG small-molecule on SG stability and structure. Based on previous experimental assays^16,71^, we utilize a data-driven procedure to develop models for ten SMCs expected to dissolve SGs and another ten that do not induce dissolution (see also Supplemental Information, Section S1). First, graph-based coarse-graining^72^ is used to obtain distinct representations for dissolving (DSMs) and non-dissolving small molecules (NDSMs) (Figures 4A,B); this approach uses graph-theoretic edge contractions to identify coarse-grained beads from the underlying molecular connectivity, providing a systematic and transferable procedure across all compounds. Homotypic interaction parameters were predicted using supervised machine learning trained on MPiPi amino and nucleic acid parameters using chemical descriptors from Mordred^73^ as inputs, and heterotypic parameters were obtained using Lorentz-Berthelot mixing rules^74,75^. The descriptors used for predictions as well as the resulting values are provided in Supplementary Information, Tables S1 and S2.

**Figure 4.**
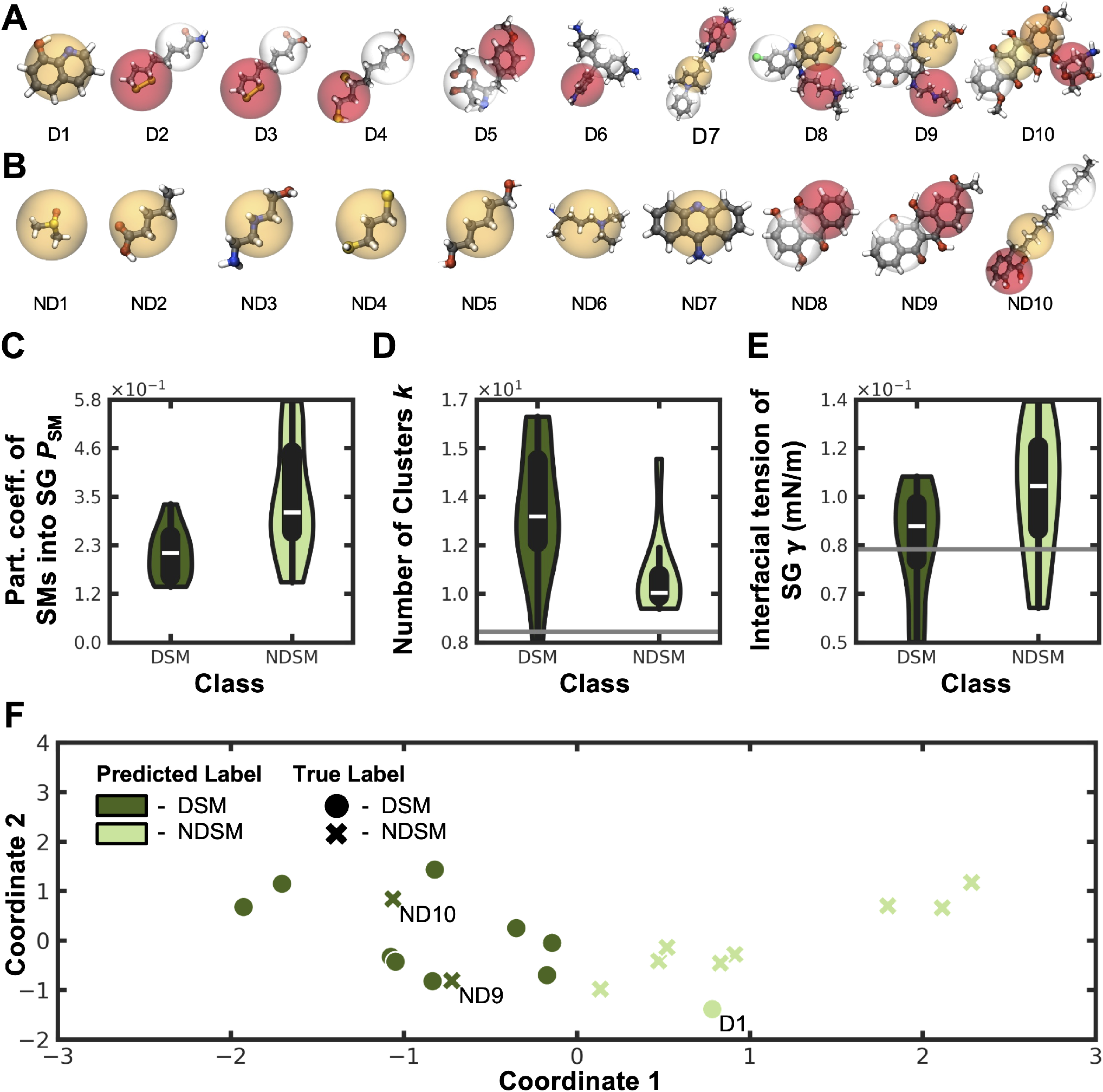
Simulations reveal distinct effects of dissolving and non-dissolving small-molecule compounds (SMCs) on SG properties. **(A, B)** In ascending order of molar mass, chemical structures and coarse-grained representations for the set of ten SMCs **(A)** with expected propensity to dissolve stress granules (D1 D10) and **(B)** without expected propensity to dissolve stress granules (ND1 - ND10). **(C-E)** Violin plot comparisons of the highest inter-class variance average measurements showcasing behavior of systems containing DSMs and NDSMs for **(C)** the partition coefficient of the small-molecule into the SG *P*_SM_, **(D)** the number of separate biopolymer clusters in the system *k*, and **(E)** the interfacial tension of the SG *γ* (mN*/*m). The horizontal gray line is a reference to the average properties of the SG control system. The violin plots use kernel density estimation for the distributions but do not extend past the extreme data points. (**F**) Classification of SMC effects on SGs based on clustering of the principal components of the small-molecule partition coefficient *P*_SM_ and the number of biopolymer clusters *k*. D1, ND9, and ND10 are noted as compounds whose assigned class differs from the experimental determination.

Overall, this approach, which avoids intensive parameterization procedures^21,76^, showcased strong capabilities to describe the MPiPi force field and produced distinct interaction potentials across the set of SMCs considered (Supplementary Information, Figure S1), allowing us to probe, at least qualitatively, the behavior of these SMCs using MD simulations. As a means of indirect validation, there is strong alignment with parameters produced by our random forest model and those obtained using more expansive, intensive procedures^77^ (Supplementary Information, Figure S1). Furthermore, control simulations of each small molecule showed no aggregation, indicating that the polarity descriptors embedded in the CG parameters yield realistic solubilities. Altogether, this highlights our approach as a reasonable means to obtain parameters for SMCs that are in qualitative alignment with chemical expectations.

Figures 4C-E illustrate that DSMs and NDSMs have distinct effects on various SG properties. Specifically, DSMs reduce the fraction of biopolymers in the largest cluster compared to both NDSMs and the SG control, suggesting that DSMs strip biopolymers from the cluster and induce destabilization and partial dissolution of the SG. This is also clear when examining the proportion of each biopolymer species that remain part of the SG (Supplementary Information, Figure S2). As a result, the number of contacts between biopolymers is higher for NDSMs than for DSMs (Supplementary Information, Figure S4), owing to the fact that biopolymers are no longer as localized in the DSM-containing systems. Numerous other properties also display distinct responses between DSMs and NDSMs (see also Figure S5 in the Supplementary Information). Perhaps counterintuitively, the partition coefficient of the DSMs is lower than that of the NDSMs (Figure 4C), which we hypothesize is due to the removal of biopolymers that interact most strongly with DSMs.

DSMs also fragment the assembly into a larger number of distinct biopolymer clusters *k* and lower the interfacial tension *γ* of the residual condensate relative to NDSMs (Figures 4D,E), consistent with a looser, more readily dispersed structure. The Green–Kubo viscosity of the condensate, by contrast, is not reduced by DSMs. At 300 K it is comparable to the control within uncertainty (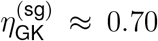 Pa s for DSMs and 0.64 Pa s for NDSMs, versus 0.56 Pa s for the control), so the simulations do not provide evidence of DSM-induced fluidization. Through the Stokes–Einstein relation evaluated with 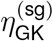 and the chain-averaged hydrodynamic radius *R*_h_, the Green–Kubo viscosity additionally provides an estimate of the intra-condensate diffusion coefficient. At 300 K, *D*^(sg)^ ≈ 0.14 µm^2^/s for the control and ≈ 0.11 and 0.09 µm^2^/s for the DSM- and NDSM-containing systems, comparable to the control within uncertainty and, like the viscosity, showing no DSM-induced enhancement. These values are of the same order as protein diffusion coefficients measured inside biomolecular condensates (*D* ≈ 0.01–0.07 µm^2^/s for PGL-3 and LAF-1 droplets)^78–80^. Likewise, the Green–Kubo viscosity (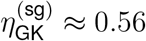 Pa s) lies at the fluid end of the range reported for protein condensates, which extends to ≈ 34 Pa s for LAF-1 droplets at physiological salt^79^ and to still higher, age-dependent values as condensates evolve into soft, glass-like Maxwell fluids^81^. As expected for coarse-grained force fields that do not explicitly target dynamical properties, these absolute values remain approximate, while relative trends across systems are preserved^21,27,82^.

Figure 4F demonstrates that examining how SMCs affect simulated SG properties can classify DSMs and NDSMs with useful, but imperfect, agreement. We perform standard scaling by removing the mean and scaling to unit variance on the two measured properties that forward feature selection identified as most discriminating between the DSM and NDSM classes: the small-molecule partition coefficient *P*_SM_ and the number of biopolymer clusters *k*. We then apply principal component analysis (PCA) to these scaled features for visualization and clustering analysis. This results in substantial separation of DSMs (dark green) and NDSMs (light green), with the associated PCA eigenvalues (Supplementary Information, Figure S8). Then using *K*-means clustering to assign labels achieves a classification accuracy of 85% (17 of 20 compounds). Because we fix this two-feature set rather than reselecting features within each fold, leave-one-out cross-validation gives the same 85% accuracy, which is significant at *p* = 0.005 against a permutation null over the class labels. On their own, *P*_SM_ and *k* classify at 65% and 75%, and a larger five-feature set (*R*^(sg)^, *P*_SM_, 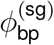, *D, η*_GK_) reaches only 70%, so the combination of *P*_SM_ and *k* separates the classes better than either feature alone or the larger set. The three misclassified compounds (D1, ND9, and ND10) sit nearest the boundary between the two clusters, and ND10 is a known counter-screen failure (see below). We suspect that, while any individual observable may exhibit overlap between DSM and NDSM populations, the net effect of observing possibly minor differences across multiple properties enables the 85% classification accuracy relative to experimental labels and reflects separation by the joint multi-feature pattern rather than any single metric.

Several results are also striking, given the approximate nature of the modeling. For example, most DSMs of lower molar mass, represented by a single CG bead, are classified correctly, as is NDSM-8, a chemical fragment of DSM-9. Some misclassifications, such as NDSM-10, are understandable, as NDSM-10 was initially identified as a DSM in an experimental screen but failed a counter-screen^16^, making it a challenging test case. Extracting properties from the MD simulations is also essential for accurate classification, as using only SMC homotypic interaction model parameters yielded a 55% classification accuracy, with an 80% false positive rate and 10% false negative rate. This highlights that the simulations, by capturing the physics of biomolecular condensates and interactions with SMCs, align well with experimental outcomes in terms of the apparent destabilization of SGs by DSMs relative to NDSMs.

### Small-molecule compounds alter internal SG structure

To further assess the differences in SG properties between DSMs and NDSMs, we compare their radial density profiles. Figures 5A,B show that DSMs reduce condensate radius relative to NDSMs (from ≈ 138 Å for the control SG and 131 Å for NDSMs to 119 Å for DSMs); NDSMs do not markedly affect the size of the condensate. Resolved by biopolymer species (Supplementary Information, Figure S2C,D), both DSMs and NDSMs affect biopolymer localization, with DSMs having a stronger effect. In the presence of DSMs (Supplementary Information, Figure S2C), there are notable changes to biopolymer distributions. TDP43 mixes into outer layers and is depleted in the center, while FUS, TIA1, PABP1, and RNA shift inward. TTP remains similar to the control, with more compact and mixed biopolymer profiles. In the presence of NDSMs (Supplementary Information, Figure S2D), effects are directionally similar but less striking than DSMs, with results being closer to the control SG. This suggests NDSMs do not significantly alter the multiphasic structure of the SG but do induce TDP43 mixing into other layers. Relative to the control SG, the positions of the midpoint concentrations for the biopolymers are more tightly grouped with DSMs than for NDSMs, and the shifts are smaller for the latter. These rearrangements are directly conveyed in Figure 5C, which shows the domain-resolved contact difference between the DSM and NDSM classes, and Figure 5D, which quantifies the accompanying change in core composition. This reveals that, relative to NDSMs, DSMs deplete G3BP1 from the condensate core (by ≈ 2 percentage points in core mass fraction) while enriching PABP1 and the coding (UCG) RNA nucleotides.

**Figure 5.**
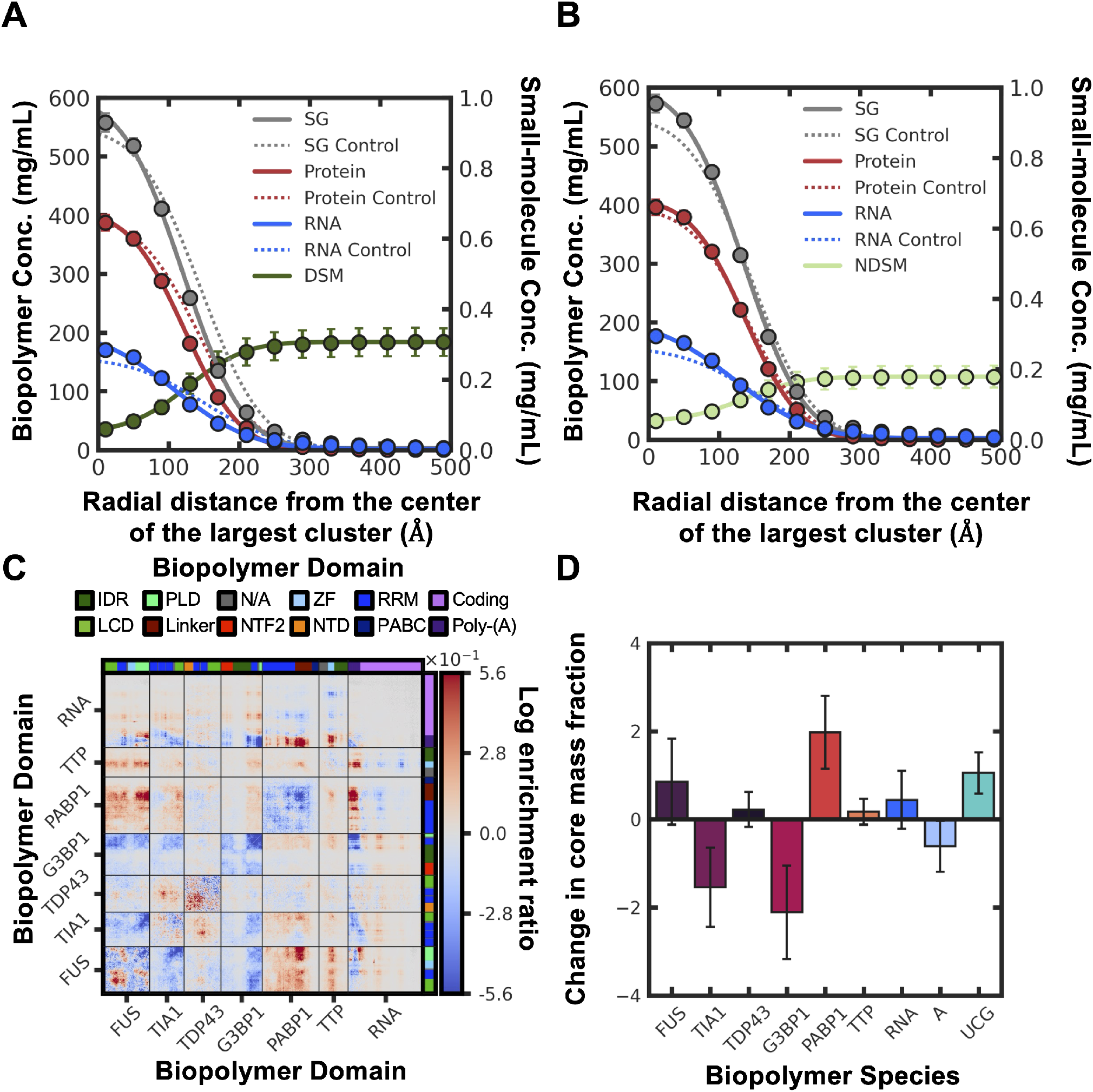
Dissolving and non-dissolving small molecules have distinct effects on biopolymer distribution in the SG. **(A)** Radial mass-density profile of SG systems containing dissolving small molecules (DSMs), resolved by biopolymer class. Solid curves are the DSM-class condensate, dotted curves the SG control; the mean DSM concentration is overlaid on the right-hand axis. **(B)** As in (A) for non-dissolving small molecules (NDSMs), with the NDSM concentration on the right-hand axis. **(C)** Domain-resolved difference contact map (DSM-class minus NDSM-class): color gives the change in log contact enrichment (log enrichment ratio) of each domain pair between the two classes. Diverging scale centered at 0 (red = enriched under DSMs, blue = enriched under NDSMs); clipped at the 99th percentile of | value | so a few extreme cells do not saturate the map. **(D)** Per-species change in core composition (DSM-class minus NDSM-class): the difference in each species’ fractional contribution to the biopolymer density within the condensate core. Bars use the per-species colors of the radial profiles; error bars are the across-compound standard error (10 DSMs, 10 NDSMs).

Overall, these results suggest that DSMs primarily interact at the SG boundary, associating with RNA in the diffuse shell and consequently stripping away the RNA scaffold. This collapses the multiphasic architecture and leaves a denser, smaller SG that resists further partitioning. Consistent with this interpretation, the fraction of each biopolymer present in the condensed phase is significantly reduced in the presence of DSMs compared to NDSMs (Supplementary Information, Figure S2). Examining concentration differences relative to the SG control confirms these trends (Supplementary Information, Figure S3), with depletion of material at the SG boundary for DSMs but a smaller impact by NDSMs. This boundary depletion is the most pronounced difference between the two classes, since both seemingly induce some enrichment in biopolymer concentration within 100 Å of the SG center. When resolved by species, however, DSMs also distinctly deplete G3BP1 throughout the SG, whereas NDSMs do not affect its distribution. Because G3BP1 is the most numerous species in the model SG and carries a strong contact signal (Figure 3A), its selective depletion by DSMs is notable. The residual compact core could in principle be more susceptible to liquid-to-solid transitions. However, the Green–Kubo viscosity of the condensate under DSMs is comparable to that of the control rather than reduced (Supplementary Information, Figure S5E), so our early-time simulations neither demonstrate nor exclude such a transition.

### Small-molecule compounds alter SG component interactions

To elucidate the disparity in structural effects between DSMs and NDSMs, we contrast how biopolymers interact based on the difference between contact probability maps obtained for DSMs and for NDSMs, in the same fashion as Figure 3.

At the biopolymer level, Figure 6A shows that DSMs increase contacts with FUS and, to a lesser extent, PABP1, while reducing interactions involving G3BP1, TIA1, and TTP. Notably, DSMs also decrease RNA contact with G3BP1 and TIA1, both of which are central to the SG structure and rely on RNA for phase separation. This supports the hypothesis that DSMs disrupt the function of RNA as a scaffold within the SG network. TDP43 contact probabilities remain largely unchanged. However, when normalized by biopolymer abundance (Figure 6B), DSMs seemingly enhance overall interaction affinity across all biopolymers. This likely reflects the fact that biopolymers remaining after DSM action are more tightly bound within the SG. The increase in normalized PABP1 contacts further suggests tighter binding, consistent with its shift toward the SG center under DSM influence (Supplementary Information, Figure S2C,D).

**Figure 6.**
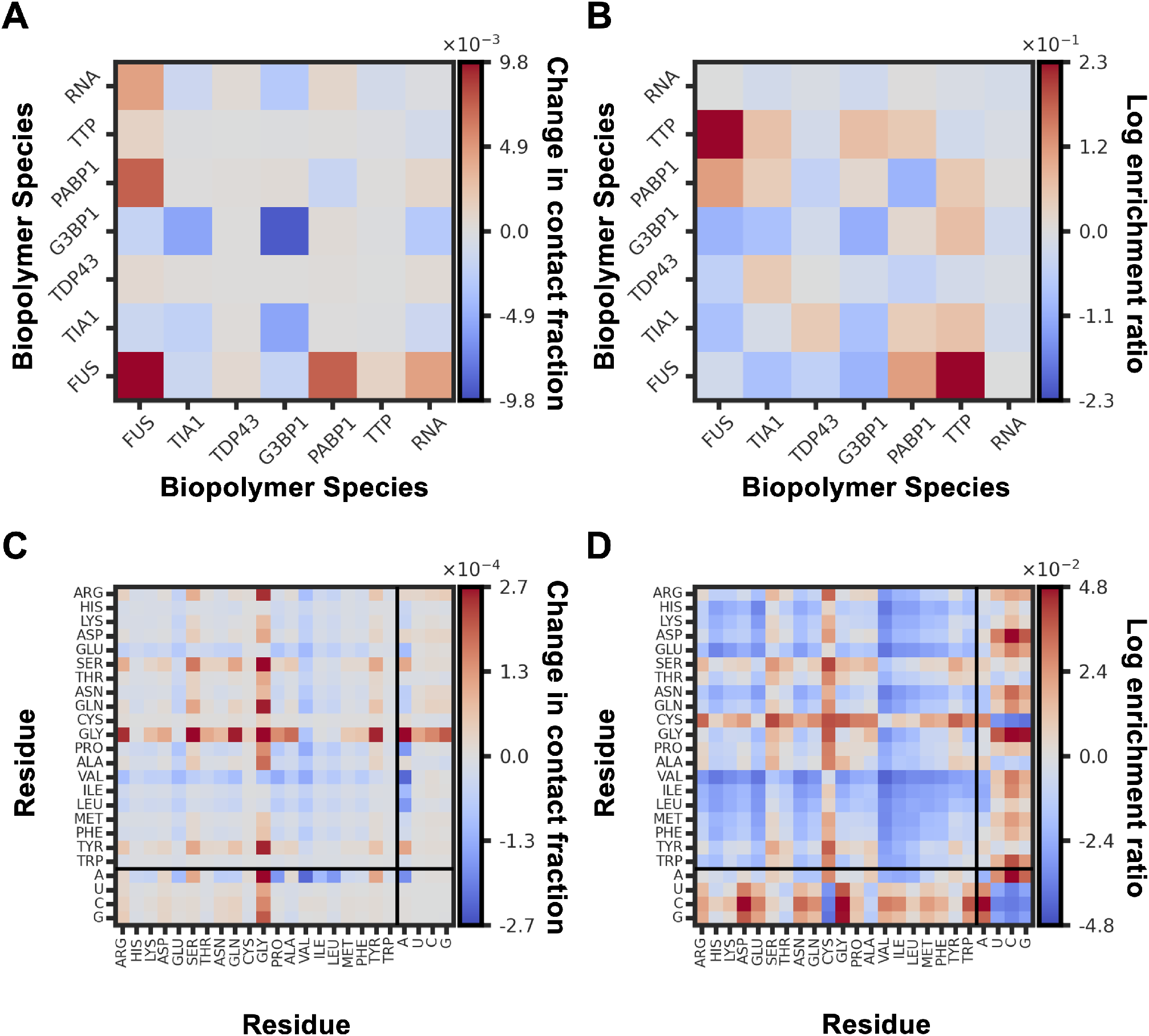
Dissolving and non-dissolving small molecules have distinct effects on biopolymer interactions. Every panel is a difference map (DSM-class minus NDSM-class condensate); inter-chain contacts only. **(A)** Change in contact fraction between biopolymer species, Δ*f* = *f*_DSM_ − *f*_NDSM_. Diverging scale centered at 0: red = the pair makes more contacts under DSMs, blue = more under NDSMs. **(B)** Change in log enrichment (log enrichment ratio) between biopolymer species, using the chain-pair null model of Figure 3(B). **(C)** Change in contact fraction between residues (amino/nucleic acids). **(D)** Change in log enrichment between residues, using the residue-abundance null model of Figure 3(D). All color scales are clipped at the 99th percentile of |value| to keep a few extreme cells from saturating the map.

Residue-resolved interaction analysis (Figure 6C) reveals a marked increase in Gly interactions with most amino acids and all nucleic acids. This is especially pronounced in FUS, which shows significantly elevated contact probabilities. After normalizing for amino and nucleic acid content (Figure 6D), we observe a general increase in interactions, supporting the hypothesis that the residual SG after partial dissolution consists of tightly bound biopolymers. Notably, several interactions associated with phase-separation, such as A-protein and RNA-RNA contacts, are enhanced. Altogether, these results suggest that DSMs selectively disrupt weaker interactions within the SG, leaving behind a more compact, strongly interacting group of biopolymers.

### Dissolving compounds preferentially interact with RNA

Compared to NDSMs, DSMs display higher affinity for RNA and interact less with most other biopolymers (Figure 7A). This aligns with earlier findings that DSMs primarily act at the SG periphery, stripping biopolymers but not penetrating the core, unlike NDSMs. When normalized by biopolymer number (Figure 7B), DSMs exhibit stronger but more varied interactions with biopolymers compared to NDSMs, with the highest affinity for RNA followed by TDP43. The positive differences in Figure 6B suggest that NDSMs behave more like inert penetrants than major disruptors of SG structure. Meanwhile, the strong DSM-RNA interactions support the hypothesis that DSMs disrupt the SG by preferentially associating with nucleic acids, weakening mRNA contacts that scaffold the SG. We emphasize that the coarse-grained model represents this preferential RNA association only through effective, isotropic pair interactions: it contains no explicit base-stacking geometry and the RNA is modeled without base-pairing, so “intercalation” is invoked as an interpretation consistent with the known aromatic, nucleic-acid-binding chemistry of these compounds rather than as a directly resolved mechanism. The small-molecule beads also carry no formal charge, so the modeled RNA preference arises from effective size/polarity (van der Waals) interactions rather than explicit electrostatics or cation–*π* contacts. The interaction with TDP43 is notable, given its role in SG aging and pathologies, and is consistent with reports that aromatic compounds prevent TDP43 aggregation in SGs^16^. At the level of amino and nucleic acids (Figures 7C,D), DSMs primarily interact with nucleic acids, especially RNA, and less frequently with Arg, Gly, Phe, Tyr, and Trp. The residues Gly and Arg are abundant in FUS, PABP1, and G3BP1, which are weaker targets of the modeled DSMs (Figure 7B). Additionally, DSMs show lower affinity for ‘A’ compared to other nucleic acids, suggesting preferential interaction with the coding region at the SG periphery rather than internal poly-(A) tails. Overall, this indicates that DSMs act by targeting RNA and RNA-dependent proteins, which may subsequently promote SG dissolution. Nevertheless, these results should not be viewed as demonstrating or capturing the complete biophysical picture of SG dissolution. Instead, the simulations identify precursor responses that appear informative for distinguishing DSM and NDSM behavior.

**Figure 7.**
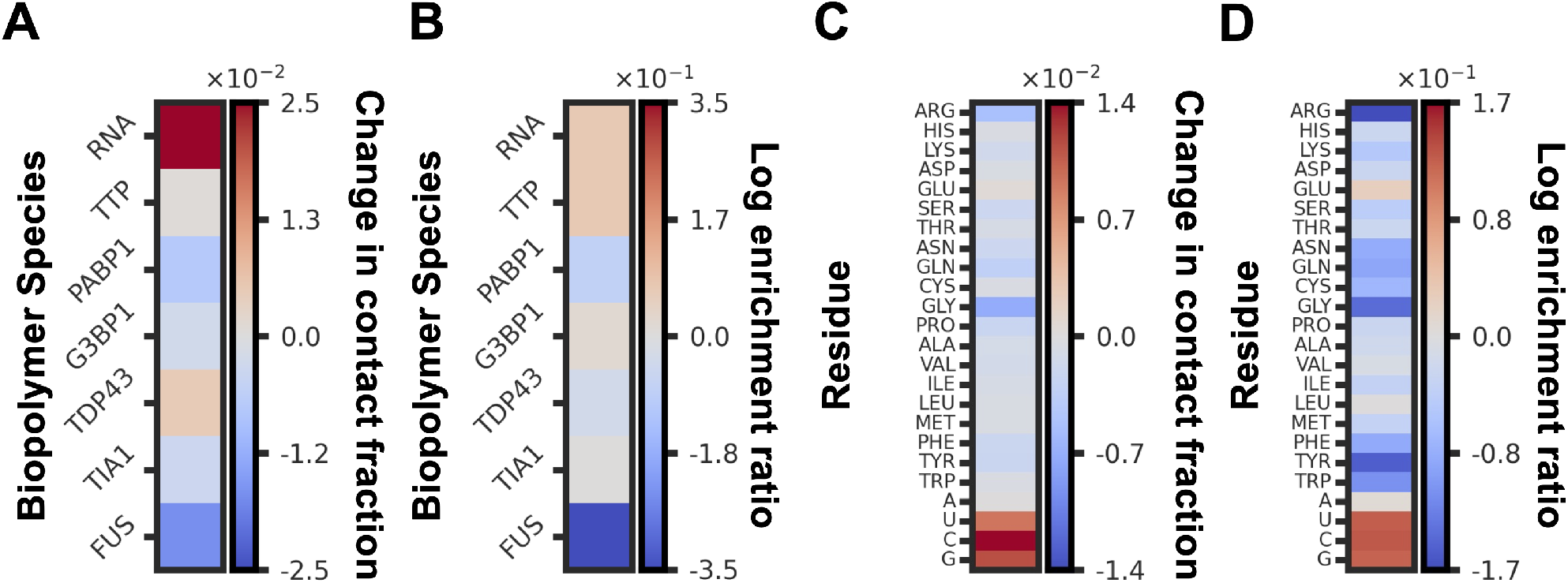
Dissolving small molecules (DSMs) preferentially interact with RNA relative to non-dissolving small molecules (NDSMs). Every panel is a difference map (DSM-class minus NDSM-class condensate) of small-molecule–biopolymer contacts; each row is the small molecule paired with one biopolymer partner. **(A)** Change in contact fraction between the small molecule and each biopolymer species, Δ*f* = *f*_DSM_ − *f*_NDSM_. Diverging scale centered at 0: red = stronger small-molecule–partner contact under DSMs. **(B)** Change in log enrichment (log enrichment ratio) between the small molecule and each biopolymer species (chain-pair null model). **(C)** Change in contact fraction between the small molecule and each residue (amino/nucleic acid). **(D)** Change in log enrichment between the small molecule and each residue (residue-abundance null model). The prominent red RNA and nucleotide bands in (A) and (B) show that DSMs concentrate their contacts on RNA. Color scales are clipped at the 99th percentile of |value|.

These computational signatures do find encouraging support in experiment. The planar, aromatic dissolving compounds modeled here were identified in a high-content screen in which such molecules dissolve stress granules by interfering with the RNA-dependent recruitment of RNAbinding proteins^16^, mirroring the RNA-scaffold disruption that emerges from our simulations. In addition, one of our dissolving compounds, lipoamide (D2), was recently shown to dissolve FUSand TDP43-associated stress granules and to ameliorate ALS phenotypes in animal models^10^; our blind, physics-based classifier independently assigns it to the dissolving class, and the experimental finding that it acts by partitioning into the condensate and altering its collective properties rather than through high-affinity binding is consistent with the partitioning-based logic of our model, in which the small-molecule partition coefficient *P*_SM_ is among the most discriminating features (Figure 4F). We caution, however, that the experimentally resolved target of lipoamide is a redox-active modification of intrinsically disordered proteins such as SFPQ, which our minimal, charge- and redox-agnostic model does not represent; the agreement therefore concerns the compound classification and the partitioning-based mode of action rather than this specific molecular chemistry^10^.

### Temperature modulates SG stability and dissolving-compound efficacy

Because both cellular stress and condensate stability are temperature dependent, we repeated the analysis across seven temperatures spanning 285–315 K (Supplementary Information, Figures S7–S10). The control SG remains assembled throughout this window but progressively loosens as temperature rises, with the dense-phase concentration decreasing, the interface broadening, and the interfacial tension proxy increasing modestly while the transfer free energy stays essentially constant. Fitting the projected binodal yields an apparent critical temperature for the control SG of *T*_*c*_ ≈ 389 K, although this value is model-dependent and the MPiPi force field is known to overestimate condensate *T*_*c*_ relative to in vitro values^83^, so we interpret only relative shifts between systems. The component proteins differ markedly in thermal robustness. FUS remains a fully embedded core anchor at every temperature, whereas TDP43 is the most thermally labile species and migrates to and beyond the interface as temperature rises, reminiscent of the surface-bound, aggregation-prone TDP43 implicated in ALS/FTD. The efficacy of the dissolving compounds also scales with temperature, as the DSM-induced reduction in the largest-cluster fraction grows roughly five-fold across the window, and even nominally non-dissolving compounds become destabilizing at the highest temperature. This temperature sensitivity manifests as a class-level signature in the apparent critical temperature, which shifts by ≈ 35 K for the DSM class and ≈ 9 K for the NDSM class relative to the control. The classifier is most accurate when its features are selected and evaluated at the same temperature, performing best at 300 K and transferring only partially across the window, such that single-temperature screening suffices for clearly-acting compounds while marginal candidates benefit from evaluation at more than one temperature.

The temperature dependence of the interaction network mirrors these structural trends. The dominant interaction motifs identified at 300 K persist across the entire window but weaken progressively as temperature increases, consistent with the falling dense-phase concentration and the broadening interface (Supplementary Information, Figures S11 and S12). These motifs include the G3BP1*/*FUS-to-RNA bridges at the species and domain level along with the Arg, Gln, Tyr, and Trp contacts and the hydrophobic Leu*/*Val*/*Ile contacts at the residue level. The G3BP1to-RNA bridge is the single most thermally sensitive contact and the one most strongly erased by DSMs, so the DSM − NDSM contrast in protein − RNA contacts is markedly larger at 315 K than at 285 K. This reinforces the picture that dissolving-compound efficacy arises from a thermally assisted disruption of the RNA-scaffold contacts that hold the multiphasic architecture together.

### Implications and Considerations

Together, these results demonstrate how a minimal multicomponent SG model can recover key features of stress-granule organization, highlight the central scaffold–surfactant role of RNA, and reveal how small molecules with distinct chemical characteristics differentially perturb this architecture. Our analyses further identify early-stage structural signatures that distinguish dissolving from non-dissolving compounds. Relative to non-dissolving compounds, dissolving compounds consistently destabilize the condensate by stripping biopolymers principally from the boundary, depleting G3BP1, and redistributing RNA–protein contacts. These signatures suggest actionable design principles for screening and optimization of SG-destabilizing compounds, such as targeting RNA-scaffold contacts or interfacial interactions. Because active cellular processes and long-timescale maturation are not modeled here, we view these simulations as a quantitative, physics-informed screen that can later be coupled to or refined with experiments to test whether similar scaffold–surfactant architectures generalize across other RNP condensates, such as Pbodies and nucleoli.

There are also notable limitations and opportunities for future work. While our SG model is substantially more complex than most computational studies of biomolecular condensates, it still falls short of the compositional and structural complexity of real SGs. Future work incorporating additional protein species, diverse RNA transcripts, and greater chemical detail will be important for developing a more complete understanding of these assemblies. In particular, the present model uses a single mRNA sequence of fixed length and nucleotide composition (a coding region with a poly-(A) tail), and both the length and the base composition of RNA are known to influence the phase behavior of RNP condensates, with shorter or compositionally distinct transcripts able to shift the balance between scaffold- and surfactant-like behavior^2,46,47^. The quantitative observations reported here may therefore be sensitive to this choice, and we expect the relative trends between dissolving and non-dissolving compounds to be more robust than the absolute values. In addition, the analysis and classification developed here rely on proxy properties that correlate with experimental dissolution rather than direct observation of complete SG disassembly. Extending simulations to longer timescales and integrating complementary *in vitro* studies will be important for connecting these early destabilization events to the full dissolution process.

The nature of the coarse-grained modeling and resolution also introduces important approximations. Folded protein domains are represented at one bead per residue and do not retain tertiary structure, so domain-specific contact patterns should be interpreted qualitatively. Likewise, RNA is modeled without explicit base pairing or base stacking, meaning that the observed poly(A)-core and coding-region-shell organization reflects composition- and charge-driven partitioning rather than sequence-specific RNA secondary structure. Finally, despite the strong alignment with experimental outcomes, the coarse-grained parameters for small molecules were obtained through an approximate data-driven procedure rather than full bottom-up validation. Establishing a more rigorous foundation for these parameters remains an important direction for future refinement.

## RESOURCE AVAILABILITY

### Lead Contact

Requests for further information and resources should be directed to and will be fulfilled by the lead contact, Michael A. Webb.

### Materials Availability

This study did not generate unique materials.

### Data and Code Availability

- All data needed to evaluate the conclusions in the paper are available in the main text or the supplementary materials.
- All code and additional (processed) data has been deposited on Github and is publicly available as of the date of publication (https://github.com/webbtheosim/stress-granule.git).
- Any additional information and other (raw) data including unprocessed molecular dynamics simulation trajectory files required to reanalyze the data reported in this paper is available from the lead contact upon request.

## ACKNOWLEDGMENTS

This work was partially supported by the National Science Foundation grant 2237470 (M.A.W). M.A.W. also gratefully acknowledges financial support from the Princeton Laboratory for Artificial Intelligence. The authors thank all members of the lab for their support. The authors thank Dr. Clifford P. Brangwynne, Dr. Anita Đonlić, Dr. Satyen Dhamankar, Dr. Jerelle Joseph, and Dr. Gerhard Hummer for helpful discussions. Authors also acknowledge assistance from Princeton Research Computing at Princeton University, which is a consortium led by the Princeton Institute for Computational Science and Engineering (PICSciE) and Office of Information Technology’s Research Computing.

## AUTHOR CONTRIBUTIONS

Conceptualization: J.L.K., M.A.W. Methodology: J.L.K., M.A.W. Investigation: J.L.K. Visualization: J.L.K., M.A.W. Writing (original draft): J.L.K., M.A.W. Writing (review and editing): J.L.K., M.A.W. Funding Acquisition: M.A.W Resources: M.A.W Supervision: M.A.W.

## DECLARATION OF INTERESTS

Authors declare that they have no competing interests.

## DECLARATION OF GENERATIVE AI AND AI-ASSISTED TECHNOLOGIES

During the preparation of this work the authors used ChatGPT-4 in order to improve readability of some sentences. After using this tool/service, the authors reviewed and edited the content as needed and take full responsibility for the content of the publication.

## SUPPLEMENTAL INFORMATION INDEX

Document SI (PDF). Figures S1–S13 and Tables S1-S3 and their legends:

Figure S1: Overview of parameterization approach for small molecule compounds.

Figure S2: Analysis of the stress granule using radial density profiles resolved by biopolymer species and a violin plot of the change in proportion of biopolymers upon exposure to DSMs and NDSMs.

Figure S3: Analysis of the stress granule using the change in the radial density profiles resolved by biopolymer class and species upon exposure to DSMs and NDSMs.

Figure S4: The difference in net contacts between systems that contain dissolving versus non-dissolving small molecule compounds, reported as the change in contact number.

Figure S5: Violin plot comparisons of the average behavior of systems containing dissolving (DSM) and non-dissolving (NDSM) small molecule compounds.

Figure S6: Additional comparison of Control, DSM, and NDSM systems.

Figure S7: Temperature dependence across *T* = 285–315 K for the control SG, DSM, and NDSM systems.

Figure S8: Robustness of the PCA + *k*-means small molecule classifier across the three screened temperatures (285, 300, and 315 K).

Figure S9: Temperature dependence of SG observables across *T* = 285–315 K for the control SG, dissolving (DSM), and non-dissolving (NDSM) small molecule systems.

Figure S10: Per-species thermal response across *T* = 285–315 K for the control SG, dissolving (DSM), and non-dissolving (NDSM) systems.

Figure S11: Inter-chain log contact enrichment maps of the control stress granule.

Figure S12: Difference (DSM class minus NDSM class) inter-chain contact maps, shown as the change in log contact enrichment (log enrichment ratio).

Figure S13: Inter-chain homotypic (self–self) contact maps for the individual protein species in the control stress granule at 300 K, shown at single-residue resolution with the axes annotated by sequence domain.

Table S1: A table of Mordred descriptors used in feature vectors for random forest models predicting different force-field parameters.

Table S2: A table of non-bonded homotypic Wang-Frenkel potential parameters for small molecule

Table S3: A table of bond-stretching parameters for small-molecules.

## STAR METHODS

### Key resources table

| REAGENT or RESOURCE | SOURCE | IDENTIFIER |
| --- | --- | --- |
| Deposited data |  |  |
| Raw | This paper | Available upon request |
| Analyzed | This paper | https://github.com/webbtheosim/stress-granule.git |
| Software and algorithms |  |  |
| LAMMPS | Thompson et al.<sup>84</sup> | https://www.lammps.org/ |
| MPiPi force field | Joseph et al.<sup>27</sup> | N/A |
| GBCG | Webb et al.<sup>72</sup> | N/A |
| Machine Learning | Pedregosa et al.<sup>85</sup> | https://scikit-learn.org/ |
| ML Coarse-Graining Procedure | This paper | https://github.com/webbtheosim/stress-granule.git |
| Analysis Scripts | This paper | https://github.com/webbtheosim/stress-granule.git |
| Other |  |  |
| Resource website for this publication | This paper | https://github.com/webbtheosim/stress-granule.git |

## Method details

### General MD simulation procedures

All MD simulations were performed using the LAMMPS^84^ simulation package and the MPiPi force-field^27^. MPiPi was selected for its complementary parameterization of RNA at the time of study. It should be noted that the recently developed CALVADOS-RNA force field^43^ also includes RNA, and a point of future interest will be assessing to what extent the choice of force field influences extraction of properties or physical insights. The MPiPi force-field uses an implicit solvent and represents each amino and nucleic acid as a single bead. CG particles interact via a combination of bonded and non-bonded energy terms. Electrostatic interactions were computed using a Coulomb term with Debye–Hückel screening for charged pairs within 35 Å, with a relative dielectric constant of 80 and a Debye screening length of 7.95 Å, corresponding to a monovalent salt concentration of 0.15 M. Short-range non-bonded interactions were modeled using a Wang–Frenkel potential^86^, with interactions beyond 25 Å neglected. The equations of motion were integrated using the velocity-Verlet algorithm with a 20 fs time step, and neighbor lists were updated every 5 steps. Unless otherwise specified, simulations were performed in the canonical ensemble at *T* = 300 K within a cubic simulation cell with fixed side lengths and periodic boundary conditions. Each dimension of the cell was set to 0.24 µm (2400 Å) to minimize finite-size effects. Temperature was controlled using the Nosé-Hoover thermostat.

### System preparation

The mRNA transcript and protein sequences were obtained from the UniProt database and converted into PDB files. Initial SG system preparation followed five steps. First, initial configurations for each biopolymer were generated by simulating isolated chains (without periodic boundary conditions) for 100 ns. The number of chains was selected to ensure the final system contained no more than 70,000 CG particles. Precise specifications are provided in Table 2. In Figure 1C-D we ran simulations over a range of protein mass fractions, defined as *w*_p_ = *m*_protein_*/m*_total_ where *m*_total_ = *m*_protein_ + *m*_RNA_. Second, the final configurations from the previous step were randomly positioned and oriented within a cubic simulation cell. Third, the system was relaxed via a 2 ns microcanonical ensemble simulation, with a maximum displacement of 0.1 Å per particle per time step. Fourth, to promote condensate formation, biopolymer chains beyond 800 Å (0.08 µm) from the center of the simulation cell experienced a weak drag force of 0.2 kcal/(mol Å) for 2 ns; this procedure was used to mitigate long diffusion times for the chains to coalesce. We note that the concentration profiles in Figure 2B-D indicate a relatively compact structure for the SG, indicating that the SG formation arises more from subsequent simulation than this initial co-localization. Fifth, the system was then simulated for 160 ns, with the final 120 ns used to assess stability based on the fraction of biopolymers in the largest cluster (Figure 1C). This procedure was equivalently applied for systems with protein mass fractions ranging from *w*_p_ = 0.0 to *w*_p_ = 1.0 in increments of 0.1, as well as for individual protein species alone (*w*_p_ = 1.0) and for individual species with RNA at *w*_p_ = 0.5.

For simulations with small molecules, 8, 324 molecules were added to the previously prepared system containing all biopolymer species at *w*_p_ = 0.5, yielding a concentration of 1 mM. The molecules were randomly inserted into the simulation cell in a region outside of the largest biopolymer cluster. In particular, the allowed insertion regions extended from the outside of a sphere with a radius of 600 Å (0.06 µm) centered in the simulation cell to the sides of the simulation cell. Systems were then simulated for 2 ns in the microcanonical ensemble, with the maximum displacement capped at 0.1 Å per particle per time step to relax configurations. This setup allows the small molecules to diffuse to the model SG for interaction as they would *in vivo* rather than spontaneously appearing within the condensate.

### Calculation of properties

From the initial configurations, systems were simulated for 2 µs to calculate properties related to SG stability, structure, and dynamics. The specific properties, described in detail below and summarized in Table 1, include radial density profiles, the number of biopolymer clusters in the system, the fraction of biopolymers present in the model SG, the radius of gyration and diameter of the model SG, the small-molecule partition coefficient, interfacial tension, the transfer free energy, viscosity of the model SG, and contact maps. For the model SG system alone, properties were mostly stable after an initial transient period; the first 50 ns is discarded as equilibration, and residual inter-segment correlation is accounted for by the block analysis described under “Quantification and statistical analysis”. In contrast, systems with small molecules, particularly DSMs, were more varied as the molecules diffused and interacted with SG components. Therefore, properties from these simulations do not represent equilibrium values, though that does not preclude use for distinguishing between different behaviors (i.e., dissolving versus non-dissolving).

**Table 1.** A table with a summary and nomenclature of calculated stress granule properties.

| Property | Symbol |
| --- | --- |
| mass density profile | $c(r)$ |
| number of biopolymer clusters | $k$ |
| fraction of biopolymers in SG | $\phi_{\text{bp}}^{(\text{sg})}$ |
| radius of the SG | $R^{(\text{sg})}$ |
| interfacial width of the SG | $W^{(\text{sg})}$ |
| partition coeff. of small-molecules into SG | $P_{\text{SM}}$ |
| interfacial tension of SG | $\gamma$ |
| transfer free energy of SG | $\Delta G_{\text{trans}}$ |
| radius of gyration of SG components | $R_g^{(\text{sg})}$ |
| percolation coeff. of SG | $\sigma$ |
| Green-Kubo viscosity in SG | $\eta_{\text{GK}}^{(\text{sg})}$ |
| contact fraction and log enrichment | $f_{ab}, E_{ab}$ |

The largest cluster of biopolymers in the system is associated with the model SG. To identify clusters, we employed^20,87,88^, where clusters are merged if any residue of a biopolymer in one cluster is within a distance *r*_c_ of any residue in another cluster. Specifically, clusters at time *t* are the connected components of the graph where biopolymers *i, j* are linked if 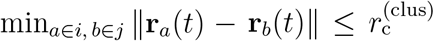 with 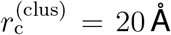 (a coarser threshold than the 16 Å contact cutoff used for the contact maps below). We then define the number of biopolymers in each cluster as 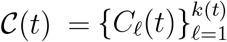. By tracking clusters throughout the trajectory, the number of unique clusters defined as *k*(*t*) = |C(*t*)|, the fraction of total biopolymers in the largest cluster, 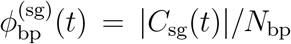 with *C*_sg_(*t*) = arg max_*C*∈*C*(*t*)_ |*C*|, and the radius of gyration of the largest cluster as 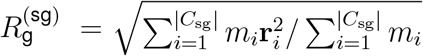.

The SG structure was characterized using radial mass density profiles (RDPs) of biopolymers associated with the largest cluster^20,89^. These were represented through *c*(*r*), the concentration (mg/mL) as function of *r*, the distance from the center-of-mass of the cluster. The profile was constructed as a mass-weighted radial histogram, with each bin accumulating the exact atomic masses of its beads, and converted from Da/Å^3^ to mg/mL by the factor 1660.539. Data were fitted to a sigmoidal function

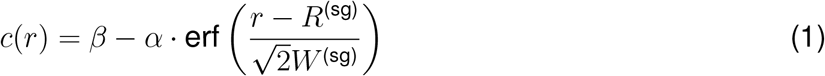

where *β, α, R*^(sg)^, and *W* ^(sg)^ are fitting parameters. The spatial extent of the SG is associated with *R*^(sg)^, while *W* ^(sg)^ is its interfacial width. The concentrations in the dense (associated with the SG) and dilute phases were obtained by *c*^(sg)^ = *β* + *α*, and *c*^(dil)^ = *β* − *α*; these fit-based concentrations agree with a cluster-average estimate given by the largest-cluster mass divided by the volume of a sphere of radius *R*^(sg)^ + *W* ^(sg)^ (e.g. ≈ 464 mg/mL for the control SG at 300 K). A quantity related to percolation was computed via

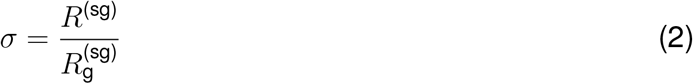

where 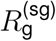 is the radius of gyration for all components in the simulation cell.

Using the concentration profiles, partitioning coefficients, for both biopolymers and small molecules, were computed by considering the ratio of component concentrations between dense and dilute phases using

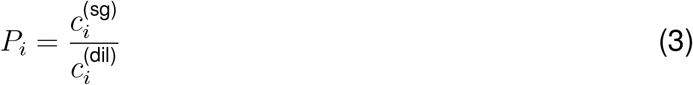

where *i* indicates the species (or set of species) included in the RDP. The free energy of transferring biopolymers from the dilute to the dense phase was estimated from the phase concentrations as Δ*G*_trans_ = *RT* ln *c*^(dil)^*/c*^(sg)^ .

Quantities related to interfacial tension^20,90^ were computed as

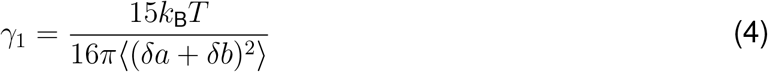

and

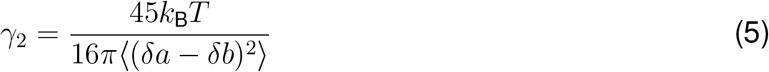

where *δa* = (*a* − *R*) and *δb* = (*b* − *R*) with *a* and *b* the axis lengths determined from principal component analysis of the mass distribution obtained as described in Ref.^20^. The mean of Eqs. (4) and (5) was used as a proxy for the interfacial tension, *γ*, for analysis. For the control SG, 134 biopolymer chains were present in total (113 protein chains and 21 RNA chains; Table 2), and 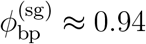 places approximately 126 chains in the condensed phase, satisfying the ≥ 100-chain criterion recommended for this shape-fluctuation estimate^20^.

**Table 2.** Biopolymer composition and concentrations of the final simulated SG system.

| Biopolymer | Residues per chain | Chains in system | Bulk concentration (mg/mL) | Mass fraction |
| --- | --- | --- | --- | --- |
| G3BP1 | 466 aa | 33 | 0.207 | 0.15 |
| TDP-43 | 414 aa | 16 | 0.086 | 0.06 |
| FUS | 526 aa | 16 | 0.103 | 0.08 |
| PABP1 | 636 aa | 16 | 0.136 | 0.10 |
| TIA1 | 386 aa | 16 | 0.082 | 0.06 |
| TTP | 326 aa | 16 | 0.065 | 0.05 |
| RNA | 840 nt | 21 | 0.684 | 0.50 |

The viscosity of the model SG was computed from the Green-Kubo relationship

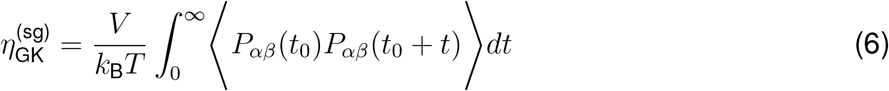

where *P*_*αβ*_(*t*) is an off-diagonal element of the pressure tensor, including contributions from all particles assigned to the largest cluster. Errors were estimated using bootstrap resampling with trajectory segments of 20 ns for computing the autocorrelation function. It is worth noting that while the right-hand side of Eq. (6) is well-defined in the CG simulations, it does not directly correspond to the physical viscosity of a condensate due to the absence of explicit solvent degrees of freedom^91^. Owing to the absence of explicit solvent, these estimates are not expected to reproduce experimental viscosities in absolute terms and are interpreted comparatively across systems; the Green–Kubo value for the control SG (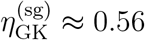 Pa s) nonetheless falls within the range reported for biomolecular condensates.

An effective, chain-averaged diffusion coefficient within the SG, *D*^(sg)^, was obtained from this Green–Kubo viscosity through the Stokes–Einstein relation

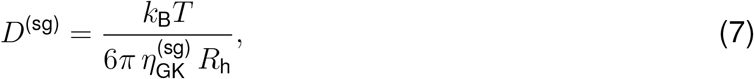

where *R*_h_ is the Kirkwood hydrodynamic radius averaged over the condensate chains. The per-chain value is computed as 1*/R*_h_ = (2*/N* ^2^) _*i<j*_ 1*/r*_*ij*_ over the *N* beads of a chain. The hydrodynamic radius is the physically appropriate length scale for this relation. Because *R*_h_*/R*_*g*_ ≈ 0.6–0.8 for flexible chains, substituting *R*_*g*_ would rescale the estimate by a comparable factor. As with the viscosity, *D*^(sg)^ is interpreted comparatively across systems rather than as an absolute physical diffusivity.

The connectivity in the SG was analyzed through inter-chain contact maps resolved at three levels, by biopolymer species, by sequence domain, and by amino/nucleic-acid type. Intra-chain contacts were excluded throughout, so sequence-local bonded neighbors within a chain do not contribute. A contact between two residues on different chains was counted when their separation was within *r*_*c*_ = 16 Å. This cutoff corresponds to approximately 2.0–2.6 *σ* across MPiPi bead types and accommodates the large nucleotide beads, for which *σ* ≈ 8.3 Å. Let *C*_*ab*_ denote the total number of inter-chain contacts between groups *a* and *b*. From this quantity, we report two complementary measures. The first is the *contact fraction*

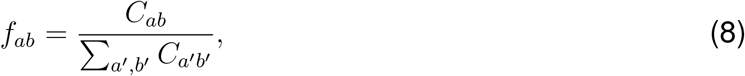

which is the share of all inter-chain contacts carried by the pair (*a, b*). The second is the *log contact enrichment*

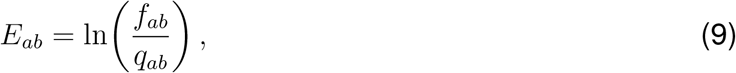

where *q*_*ab*_ is the contact fraction expected under a null model. For the biopolymer-species and domain-resolved maps, the null is a chain-pair null, *q*_*ab*_ ∝ *n*_*a*_*n*_*b*_, with *n* the number of chains and independent of chain length. For the amino/nucleic-acid maps, the null is a residue-abundance null, proportional to the product of each residue type’s abundance in the condensate. Positive *E*_*ab*_ indicates contacts more frequent than expected by chance, and negative *E*_*ab*_ indicates contacts less frequent than expected. Domain-resolved maps (e.g. Figure 2C) decompose each chain into its constituent sequence domains (IDR, PLD, LCD, Linker, NTF2, NTD, ZF, RRM, PABC, and the Coding/Poly-(A) RNA segments). These maps are displayed at single-residue resolution with the axes annotated by domain rather than averaged into domain blocks. All species share a single contact-fraction color scale in the pooled domain map, so the sparser self–self blocks are not individually legible. Supplementary Information, Figure S13 therefore isolates each species’ self–self block from the full inter-chain contact-fraction matrix and rescales it to its own range [0, max *f*_*ab*_], which resolves each biopolymer’s internal homotypic contact structure. Small-molecule contact maps were constructed identically, pairing each compound with each biopolymer or residue partner.

To highlight the effects of small molecules, difference maps between conditions report the change in contact fraction, 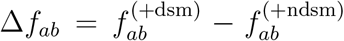, and the change in log enrichment (the “log enrichment ratio”). The color scale of difference maps was clipped at the 99th percentile of |value| so that a small number of extreme cells does not saturate the map.

### Phase behavior and apparent critical temperature

To probe thermal stability, all properties were recomputed at seven temperatures, *T* = 285, 290, 295, 300, 305, 310, and 315 K, with 300 K as the base condition. The DSM/NDSM screen was performed at 285, 300, and 315 K. From the temperature-dependent dense- and dilute-phase concentrations determined from Eq. (1), we approximated a coexistence (binodal) curve. The two branches were fitted jointly with an order-parameter law, *c*^(sg)^ − *c*^(dil)^ ∝ (*T*_*c*_ − *T* )^*β*^, with the critical exponent fixed at *β* = 0.325 and combined with a rectilinear-diameter relation for 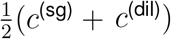, which together yield an apparent critical temperature *T*_*c*_. The simulated window (285 – 315 K) lies well below the fitted transition, and MPiPi is known to overestimate condensate critical temperatures^83^, so the absolute value is model-dependent (e.g. *T*_*c*_ ≈ 389 K for the control SG under the constrained fit) and only relative *T*_*c*_ shifts between systems are interpreted. Per-compound apparent *T*_*c*_ values were obtained from constrained fits that share the control-branch shape.

Per-species thermal localization (Supplementary Information, Figure S10) was quantified from the chain centers of mass using two measures, the condensate occupancy of each species and the mean normalized radial position *r/R*. The condensate occupancy is the fraction of analyzed frames in which a species’ chains belong to the tracked largest cluster. For the radial position, 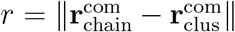 is averaged over the frames a chain is inside the condensate, and *R* is a measure of the condensate radius given by the root-mean-square radial spread of the inside chains. Evaluated per chain over inside-condensate frames, *r/R* reports where a species’ chains reside, which can differ from the peak of its density profile. Because the radial volume element grows as *r*^2^, a broadly distributed species may peak in concentration near the center yet have most of its chains at larger radii. We therefore take *r/R* as the measure of radial localization.

### Classification of small-molecule effects

Each compound was represented by a feature vector of its aggregate SG-system observables, standard-scaled to zero mean and unit variance. Discriminating features were chosen by greedy forward selection over this pool (candidates ranked by absolute Pearson correlation with the experimental label, then grown to the highest-accuracy subset), which retained two: the small-molecule partition coefficient *P*_SM_ and the number of biopolymer clusters *k*. Adding any further observable did not improve, and generally degraded, the clustering accuracy. The scaled features were projected with principal component analysis and labeled by *K*-means clustering (*K* = 2), with cluster-to-class orientation fixed on the training data. Because this two-feature set is fixed rather than reselected within each fold, leave-one-out cross-validation reproduces the in-sample accuracy exactly (85%, 17*/*20 in both cases), and its significance was established against a permutation null over the class labels (*p* = 0.005, 200 label permutations). On their own the two features classify at 65% (*P*_SM_) and 75% (*k*), while a five-feature set of structural and dynamical observables (*R*^(sg)^, *P*_SM_, 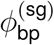, *D, η*_GK_) gives 70%.

### Small-molecule selection

To examine whether our model SG was sensitive to small molecule interactions, a set of experimentally tested compounds were selected based on their propensity to dissolve SGs^16,71,92^. In particular, twenty small molecules were chosen, evenly split between dissolving (DSM) and non-dissolving (NDSM) classes. For clarity, “DSM” and “NDSM” refer to compounds experimentally reported to dissolve or not dissolve stress granules under the assay conditions. These labels are used to select and evaluate simulation-derived classifier features, but not to parameterize the small-molecule force field or prescribe the MD response. The ten DSMs were selected from small molecules experimentally reported to dissolve SGs through direct physicochemical effects on the condensate rather than by modulating the cellular stress response^16,71^; these were labeled D1-D10. For comparison, the ten NDSMs (ND1-ND10) were selected with the following rationale. ND1 was included as it was a control in experimental studies. The set ND2, ND3, ND4, ND6, ND7, ND8, and ND9 were selected because they are chemical fragments of D4, D9, D8, and D10^71^, making them challenging test cases. ND5 (1,6-hexanediol) was included to assess whether our analyses and methods could differentiate compounds effective at low versus high concentrations, as it disrupts SG stability only at high concentrations. ND10 was included because it was identified as a hit in an initial screen but failed to inhibit SG formation in a dose-dependent manner during a counterscreen^16^.

### Small-molecule coarse-graining

The Graph-Based Coarse-Graining (GBCG) algorithm with spectral grouping was used to generate CG representations of all small-molecule compounds^72^. This spectral grouping approach successively combines nodes in a molecular graph based on eigenvector centrality rankings. Four iterations of grouping were performed, using adjacency matrices that incorporated both connectivity and node mass. To ensure no CG bead exceeded the molecular weight of the largest amino acid (Trp), nodes were merged only if the resulting mass remained below 205 Da. This procedure produces a set of topologically CG beads to represent each small-molecule at a resolution consistent with the rest of the MPiPi force field.

A data-driven parameterization approach was employed to obtain force-field parameters for the derived CG beads. The key idea is to exploit the preexisting and extensive parameterization/validation of the MPiPi force field as the basis for obtaining CG parameters for small molecules. This is achieved by using machine learning to approximate the mapping of chemical structure to coarse-grained parameters, such that CG parameters of small molecules can be inferred by similarity between chemical moieties in the small molecules to those appearing across the combination of amino and nucleic acids. To do so, each CG bead was assigned a SMILES string representing its atomistic structure and connectivity^93^, which was used to generate a feature vector of Mordred descriptors^73^. The same procedure was applied to the twenty amino acids and four nucleic acids in the MPiPi force field^27^. A random forest regression model was trained using scikit-learn^85^ to predict MPiPi homotypic parameters from these feature vectors. A greedy forward-selection method based on maximum feature variance was employed to determine a relevant subset of descriptors: each descriptor with the highest remaining variance was added to the feature set if it improved the model’s coefficient of determination (*R*^2^) on a test set; otherwise, it was discarded. This process was repeated until all descriptors were considered. A train-test split, comprising 19 amino/nucleic acids for training and 5 for testing (approximately an 80/20 split), was used to assess model performance. The results are reported in Supplementary Information, Table S1. Finally, the trained model was used to predict homotypic parameters for each CG bead of the small molecules. The parameters obtained by this approach are reported in the Supplementary Information, Table S2. Parameters for bonded interactions are reported in Supplementary Information, Table S3.

Heterotypic interaction parameters between small molecules and between small molecules and amino/nucleic acids were specified using mixing rules. Although the MPiPi force field employs explicit heterotypic parameters, this approach would be labor intensive for the small molecules. Thus, we sought to identify an appropriate mixing rule that could approximate heterotypic parameters from homotypic parameters. Four widely used mixing rules were examined: Lorentz-Berthelot^74,75^, Waldman-Hagler^94^, Fender-Halsey^95^, and Kong^96^ rules. Each rule was evaluated according to the sum of squared residuals between the explicit MPiPi heterotypic parameters and those derived from the mixing rules. The Lorentz-Berthelot rules minimized this metric, resulting in *σ*_*ij*_ and *R*_*ij*_ determined as arithmetic means and *ϵ*_*ij*_, *µ*_*ij*_, and *ν*_*ij*_ as geometric means of the homotypic parameters. The goal is qualitative, screening-level assessment of how chemical features might shift condensate organization; we therefore interpret SM effects comparatively across compounds rather than as absolute predictions.

### Quantification and statistical analysis

All statistical analyses can be found in the ‘Method details’ section of the ‘STAR Methods’ and are explained in detail here. For aggregate static and structural observables reported in the main figures, each 2 µs trajectory was split into forty 50 ns segments. The first segment was discarded (treated as additional equilibration), and the remaining segments were used to compute means and sample standard errors. The same starting configuration was used for all systems, so block uncertainties quantify uncertainty within each simulated trajectory rather than uncertainty over independent system preparations. Errors for derived quantities were obtained using error propagation. To account for temporal correlation between segments, the reported uncertainties additionally use correlation-corrected block analysis: the effective block size is chosen from a Flyvbjerg–Petersen superblock plateau combined with the PyMBAR statistical inefficiency^97,98^, the standard error is 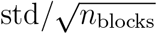 over the decorrelated blocks, and 95% confidence intervals are taken from the Student-*t* distribution. Property averages for DSMs and NDSMs were taken as the mean and standard error across the ten chemically distinct compounds in each class. These class-level SEMs therefore describe across-compound variation, not independent replicate trajectories for a single compound.

### Additional resources

Github Page: https://github.com/webbtheosim/stress-granule.git

## Notes

https://github.com/webbtheosim/stress-granule

### Competing Interest Statement

The authors have declared no competing interest.

### Summary of Updates

This revised version includes updates to clarify the modeling approach, expand validation, and better contextualize the conclusions. Major changes include new temperature-dependent simulations and projected phase-behavior analyses, expanded discussion of small-molecule parameterization and model limitations, additional domain- and contact-resolved analyses of stress granule architecture, clearer reporting of simulation system composition and protocols, and revised language distinguishing early destabilization or partial dissolution from complete stress granule disassembly.

