## Supplementary text, figures, and tables. for "Interactions Underlying Stress Granule Structure and Therapeutic Dissolution"

### Supplementary Material for Interactions Underlying Stress Granule Structure and Therapeutic Dissolution

### Contents

**S1 Data-driven coarse-graining of small molecules** **SI-2**

**S2 Force field parameters** **SI-5**

**S3 Additional analysis on small-molecule effects** **SI-7**

**S4 Temperature dependence of stress-granule structure and dissolution** **SI-14**

**S5 Per-species homotypic contact maps** **SI-18**

### S1 Data-driven coarse-graining of small molecules

To obtain coarse-grained (CG) structures of the small-molecule compounds (SMCs) tested against the stress granule (SG) model, we use the spectral graph-based coarse-graining (GBCG) algorithm of Webb and co-workers (see Methods of the main text). The spectral graph-based coarse-graining scheme produces a series of hierarchically linked molecular representations guided by spectral properties of the chemical adjacency matrix and yielding reasonable representations relative to *ad hoc* assignments. To model interactions with the SMCs, a two-step parameterization procedure is employed. The first step uses a random forest regression model to predict homotypic interaction parameters from a feature vector of chemical descriptors calculated using Mordred[1]. The model is trained on data from MPiPi, leveraging its pre-existing parameterization of amino and nucleic acids, and exploits the chemical similarity between those residues and SMC structures through an optimized descriptor-to-parameter mapping. Supplementary Information, Fig. S1 illustrates the capacity of the model to predict homotypic force-field parameters; models are trained on approximately 80% of the parameterized amino and nucleic acids (i.e., 19/24) using various feature sets and then tested on the remaining 20% (i.e., 5/24). All models perform well, with the $R^2$ values for every parameter exceeding 0.85 with a sufficiently large feature set. Additionally, in Fig. S1B we see that the DSMs tend to have less variable interaction energies than NDSMs. The second step identifies an appropriate mixing rule for handling heterotypic interactions. The use of interaction mixing rules for determining the heterotypic interactions is sensible given this difference in homotypic interaction parameters. In the MPiPi force-field, all interactions, both homotypic and heterotypic, are explicitly parameterized. To prevent extrapolation and over-reliance on the model, we avoid explicit parameterization for the SMCs and rather examined common parameter-mixing rules to determine how well they could reproduce the amino/nucleic acid heterotypic interaction parameters for MPiPi. Although no mixing rule performs universally well, the Lorentz-Berthelot mixing rules[2,3] most closely resembled the parameterized heterotypic interactions of the MPiPi force-field and were thus used for analysis.

A greedy forward-selection approach was used to identify subsets of Mordred descriptors for predicting homotypic interaction parameters via machine learning. Fig. S1 shows the random forest learning progress and final homotypic interaction parameter curves. Table S1 lists the descriptors that improved model performance on held-out amino/nucleic acids and were subsequently used to predict small-molecule parameters.

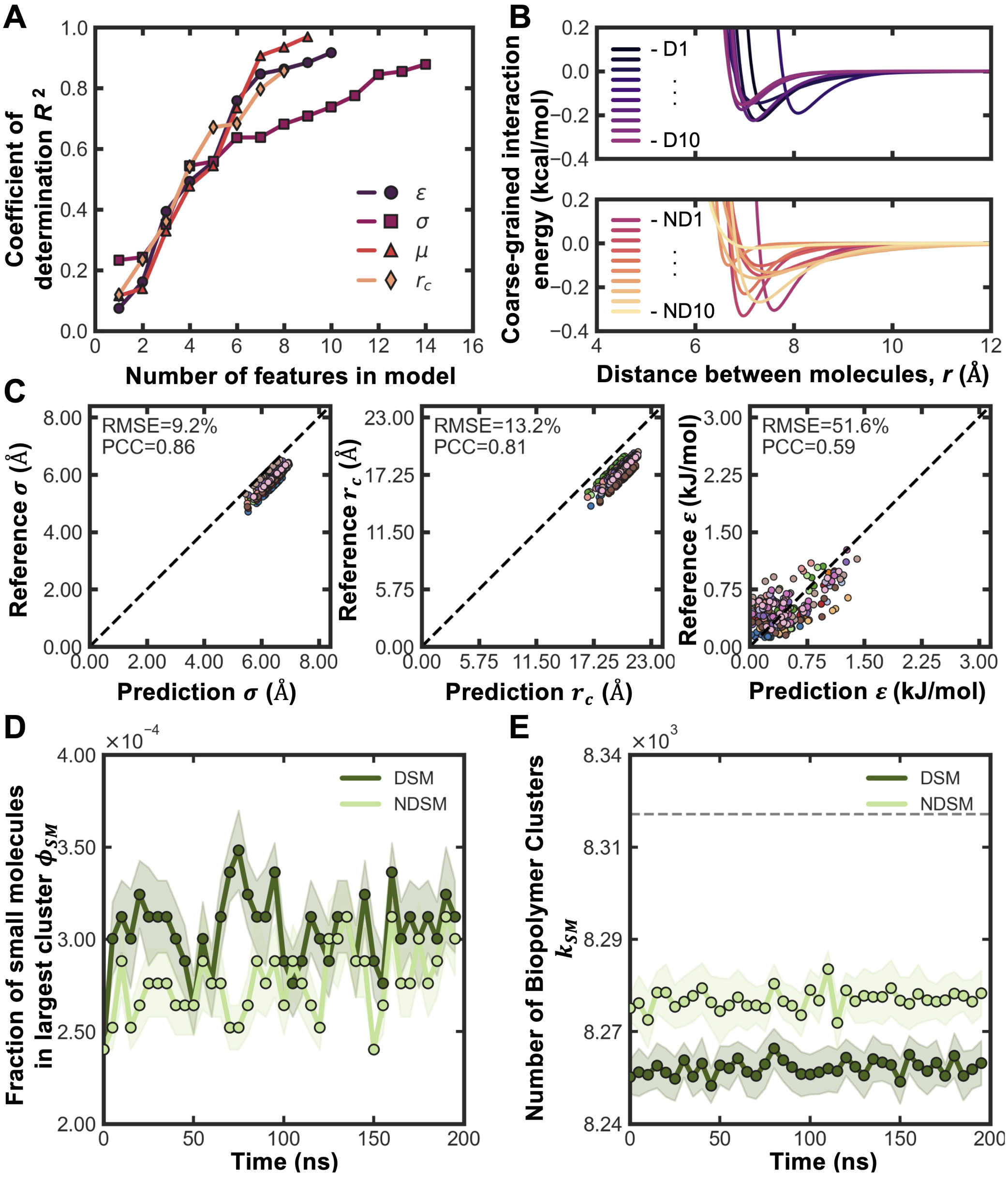


**Figure S1:** Overview of parameterization approach for small-molecule compounds. **(A)** Performance of random forest regression as a function of feature-selection number and force field parameter. Separate models are trained to predict the four different Wang-Frenkel CG interaction parameters. **(B)** Homotypic interaction curves predicted by random forest models for the small-molecule compounds split between the ten DSMs (top) and the ten NDSMs (bottom). **(C)** Parity plots of other existing small molecule and amino acid interaction parameters[4] versus our random forest regression method (colored by small molecule type). **(D)** The fraction of small molecules, $\phi_{\text{sm}}$, in the largest cluster over time for the isolated average DSM and NDSMs. **(E)** The number of small molecule clusters, $k$, over time for the isolated average DSM and NDSMs.

**Table S1:** Mordred descriptors used in feature vectors for random forest models predicting different force-field parameters.

| Parameter | MORDRED Descriptor | $R^2$ | Mean Absolute Error |
|---|---|---|---|
| $\varepsilon_{ii}$ (kcal/mol) | ATS5se | 0.074 | 0.109 |
| | WPath | 0.161 | 0.122 |
| | ATSC8v | 0.393 | 0.133 |
| | ATS6s | 0.493 | 0.119 |
| | ATS3s | 0.557 | 0.118 |
| | ATS8s | 0.758 | 0.067 |
| | ATS0se | 0.846 | 0.064 |
| | ATSC0dv | 0.863 | 0.062 |
| | Diameter | 0.884 | 0.050 |
| | Xp-3dv | 0.916 | 0.044 |
| $\sigma_{ii}$ (Å) | ATSC8v | 0.233 | 0.861 |
| | ATS5s | 0.243 | 0.774 |
| | BertzCT | 0.350 | 1.112 |
| | ATS8se | 0.544 | 0.676 |
| | ATSC5Z | 0.558 | 0.407 |
| | SpAD_Dzm | 0.637 | 0.527 |
| | SpAD_Dzse | 0.637 | 0.381 |
| | SpAbs_DzZ | 0.681 | 0.532 |
| | SpAbs_Dzare | 0.708 | 0.467 |
| | VMcGowan | 0.738 | 0.464 |
| | ATSC8Z | 0.774 | 0.309 |
| | SpDiam_Dzm | 0.845 | 0.312 |
| | MWC01 | 0.855 | 0.288 |
| | SlogP_VSA10 | 0.878 | 0.282 |
| $\mu_{ii}$ | ATS3i | 0.116 | 0.250 |
| | ATS1dv | 0.139 | 0.312 |
| | TopoPSA(NO) | 0.329 | 0.226 |
| | TIC3 | 0.477 | 0.204 |
| | SpAbs_D | 0.544 | 0.276 |
| | SpAD_DzZ | 0.734 | 0.206 |
| | SpAbs_Dzm | 0.907 | 0.100 |
| | ATSC1s | 0.935 | 0.084 |
| | ZMIC5 | 0.969 | 0.048 |
| $r_{c,ii}$ (Å) | ATSC0v | 0.119 | 1.291 |
| | SpAD_Dzpe | 0.234 | 1.997 |
| | SpAD_Dzare | 0.360 | 1.933 |
| | ATSC0dv | 0.541 | 1.961 |
| | SpDiam_Dzse | 0.671 | 1.533 |
| | ATSC6dv | 0.683 | 1.234 |
| | AATS1m | 0.796 | 1.153 |
| | LogEE_Dzse | 0.856 | 0.902 |

### S2 Force field parameters

The specific homotypic force-field parameters predicted from the ML models and used to run the coarse-grained simulations with small molecules are in Table S2. Bonded parameters were simply adapted from MPiPi (Table S3) and bond lengths from the resulting GBCG to preserve topology. Heterotypic parameters were generated directly from the Table S2 homotypic parameters using the Lorentz-Berthelot mixing rules described in the main-text Methods.

**Table S2:** Non-bonded Homotypic Wang-Frenkel Potential Parameters for Small Molecules.

| SMC ID | Name | Formula | Bead | $m$ (g/mol) | $Q$ (e) | $\varepsilon_{ii}$ (kcal/mol) | $\sigma_{ii}$ (Å) | $\nu_{ii}$ | $\mu_{ii}$ | $R_{ii}$ (Å) |
|---|---|---|---|---|---|---|---|---|---|---|
| ND1 | Dimethyl sulfoxide | $C_2H_6OS$ | 1 | 78.12922 | 0.0 | 0.100410 | 6.507146 | 1 | 8 | 18.442294 |
| ND2 | Valeric acid | $C_5H_{10}O_2$ | 1 | 102.1335 | 0.0 | 0.022258 | 6.761110 | 1 | 3 | 18.756255 |
| ND3 | Ethylenediamine | $C_2H_8N_2$ | 1 | 104.15244 | 0.0 | 0.133458 | 6.823508 | 1 | 2 | 18.740782 |
| ND4 | Propanedithiol | $C_3H_8S_2$ | 1 | 108.21676 | 0.0 | 0.269210 | 6.822550 | 1 | 3 | 19.185204 |
| ND5 | Hexanediol | $C_6H_{14}O_2$ | 1 | 118.17638 | 0.0 | 0.159666 | 6.556188 | 1 | 2 | 18.015101 |
| ND6 | N,N-diethyl-1,4-pentanediamine | $C_9H_{22}N_2$ | 1 | 158.28774 | 0.0 | 0.230895 | 6.732767 | 1 | 5 | 17.748330 |
| ND7 | 9-Aminoacridine | $C_{13}H_{10}N_2$ | 1 | 194.2361 | 0.0 | 0.332215 | 6.700928 | 1 | 5 | 24.948550 |
| ND8 | Anthraquinone | $C_{14}H_8O_2$ | 1 | 136.10708 | 0.0 | 0.180603 | 7.141808 | 1 | 2 | 23.004776 |
| | | | 2 | 104.10828 | 0.0 | 0.025928 | 6.684247 | 1 | 5 | 23.329099 |
| ND9 | 8-acetyl-6,11-dihydroxy-7,8,9,10-tetrahydro-naphthacene-5,12-dione | $C_{20}H_{16}O_5$ | 1 | 156.14068 | 0.0 | 0.293405 | 7.782794 | 1 | 5 | 24.765770 |
| | | | 2 | 180.20384 | 0.0 | 0.319659 | 6.816082 | 1 | 5 | 19.251745 |
| ND10 | Anacardic acid | $C_{22}H_{36}O_3$ | 1 | 113.22349 | 0.0 | 0.170891 | 6.980801 | 1 | 3 | 18.764813 |
| | | | 2 | 110.19958 | 0.0 | 0.170891 | 6.980801 | 1 | 3 | 18.764813 |
| | | | 3 | 125.10405 | 0.0 | 0.106535 | 6.585695 | 1 | 2 | 17.153702 |
| D1 | 8-Hydroxyquinoline | $C_9H_7NO$ | 1 | 145.16089 | 0.0 | 0.191857 | 7.766905 | 1 | 5 | 24.408175 |
| D2 | Lipoamide | $C_8H_{15}NOS_2$ | 1 | 100.1408 | 0.0 | 0.023645 | 6.756340 | 1 | 2 | 18.751684 |
| | | | 2 | 105.19285 | 0.0 | 0.271504 | 6.840142 | 1 | 8 | 18.366873 |
| D3 | Lipoic acid | $C_8H_{14}O_2S_2$ | 1 | 101.12553 | 0.0 | 0.022258 | 6.761110 | 1 | 3 | 18.756255 |
| | | | 2 | 105.19285 | 0.0 | 0.271504 | 6.840142 | 1 | 8 | 18.366873 |
| D4 | Dihydrolipoic acid | $C_8H_{16}O_2S_2$ | 1 | 101.12553 | 0.0 | 0.022258 | 6.761110 | 1 | 3 | 18.756255 |
| | | | 2 | 107.20879 | 0.0 | 0.269210 | 6.822550 | 1 | 3 | 19.185204 |
| D5 | Anisomycin | $C_{14}H_{19}NO_4$ | 1 | 158.17754 | 0.0 | 0.330791 | 7.525393 | 1 | 5 | 18.104420 |
| | | | 2 | 107.13219 | 0.0 | 0.030903 | 6.779316 | 1 | 5 | 18.261337 |
| D6 | Pararosaniline | $C_{19}H_{17}N_3$ | 1 | 195.24407 | 0.0 | 0.323070 | 6.690598 | 1 | 5 | 25.009365 |
| | | | 2 | 92.12052 | 0.0 | 0.028327 | 6.704187 | 1 | 6 | 23.061942 |
| D7 | Pyrvinium | $C_{26}H_{28}N_3$ | 1 | 91.11255 | 0.0 | 0.028327 | 6.704187 | 1 | 6 | 23.061942 |
| | | | 2 | 117.17073 | 0.0 | 0.151192 | 7.030610 | 1 | 5 | 17.896176 |
| | | | 3 | 174.24598 | 0.0 | 0.273592 | 6.759864 | 1 | 5 | 19.338650 |
| D8 | Quinacrine | $C_{23}H_{30}ClN_3O$ | 1 | 112.53861 | 0.0 | 0.177764 | 6.696324 | 1 | 5 | 22.959961 |
| | | | 2 | 106.12422 | 0.0 | 0.030903 | 6.779316 | 1 | 5 | 18.261337 |
| | | | 3 | 181.30177 | 0.0 | 0.252500 | 6.576281 | 1 | 5 | 18.522622 |
| D9 | Mitoxantrone | $C_{22}H_{28}N_4O_6$ | 1 | 188.13948 | 0.0 | 0.360487 | 7.186907 | 1 | 3 | 24.707024 |
| | | | 2 | 141.19341 | 0.0 | 0.175598 | 6.778569 | 1 | 5 | 17.589042 |
| | | | 3 | 115.15547 | 0.0 | 0.140055 | 6.589352 | 1 | 5 | 18.456455 |
| D10 | Daunorubicin | $C_{27}H_{29}NO_{10}$ | 1 | 134.13462 | 0.0 | 0.200500 | 7.314192 | 1 | 5 | 22.956225 |
| | | | 2 | 81.05077 | 0.0 | 0.031016 | 6.565698 | 1 | 5 | 19.495940 |
| | | | 3 | 139.13099 | 0.0 | 0.288206 | 6.954199 | 1 | 5 | 17.836432 |
| | | | 4 | 173.21245 | 0.0 | 0.379496 | 6.849942 | 1 | 5 | 19.510541 |

**Table S3:** Bond-stretching parameters.

| $i - j$ | $K_{ij}$ (kcal · mol$^{-1}$ · Å$^{-2}$) | $R_{ij}^{(0)}$ (Å) |
|---|---|---|
| $ND8_1$ - $ND8_2$ | 9.6 | 4.064 |
| $ND9_1$ - $ND9_2$ | 9.6 | 6.058 |
| $ND10_1$ - $ND10_2$ | 9.6 | 9.671 |
| $ND10_2$ - $ND10_3$ | 9.6 | 5.262 |
| $D2_1$ - $D2_2$ | 9.6 | 6.217 |
| $D3_1$ - $D3_2$ | 9.6 | 6.225 |
| $D4_1$ - $D4_2$ | 9.6 | 6.570 |
| $D5_1$ - $D5_2$ | 9.6 | 5.041 |
| $D6_1$ - $D6_2$ | 9.6 | 4.827 |
| $D7_1$ - $D7_2$ | 9.6 | 4.729 |
| $D7_2$ - $D7_3$ | 9.6 | 7.577 |
| $D8_1$ - $D8_2$ | 9.6 | 6.535 |
| $D8_1$ - $D8_3$ | 9.6 | 6.535 |
| $D9_1$ - $D9_2$ | 9.6 | 6.535 |
| $D9_1$ - $D9_3$ | 9.6 | 7.976 |
| $D9_2$ - $D9_3$ | 9.6 | 5.688 |
| $D10_1$ - $D10_2$ | 9.6 | 3.509 |
| $D10_2$ - $D10_3$ | 9.6 | 4.225 |
| $D10_3$ - $D10_4$ | 9.6 | 4.810 |

### S3 Additional analysis on small-molecule effects

In the control SG system, the vast majority of biopolymer species are found within the SG and distributed as shown in Fig. S2A. At a population level, we find that small molecules characterized as “dissolving” have a larger impact on the fraction of those biopolymers that remain in the SG, relative to those characterized as “non-dissolving” (Fig. S2B). The impact of dissolving and non-dissovling compounds is then shown in Fig. S2C,D, with differences more finely resolved in Fig. S3.

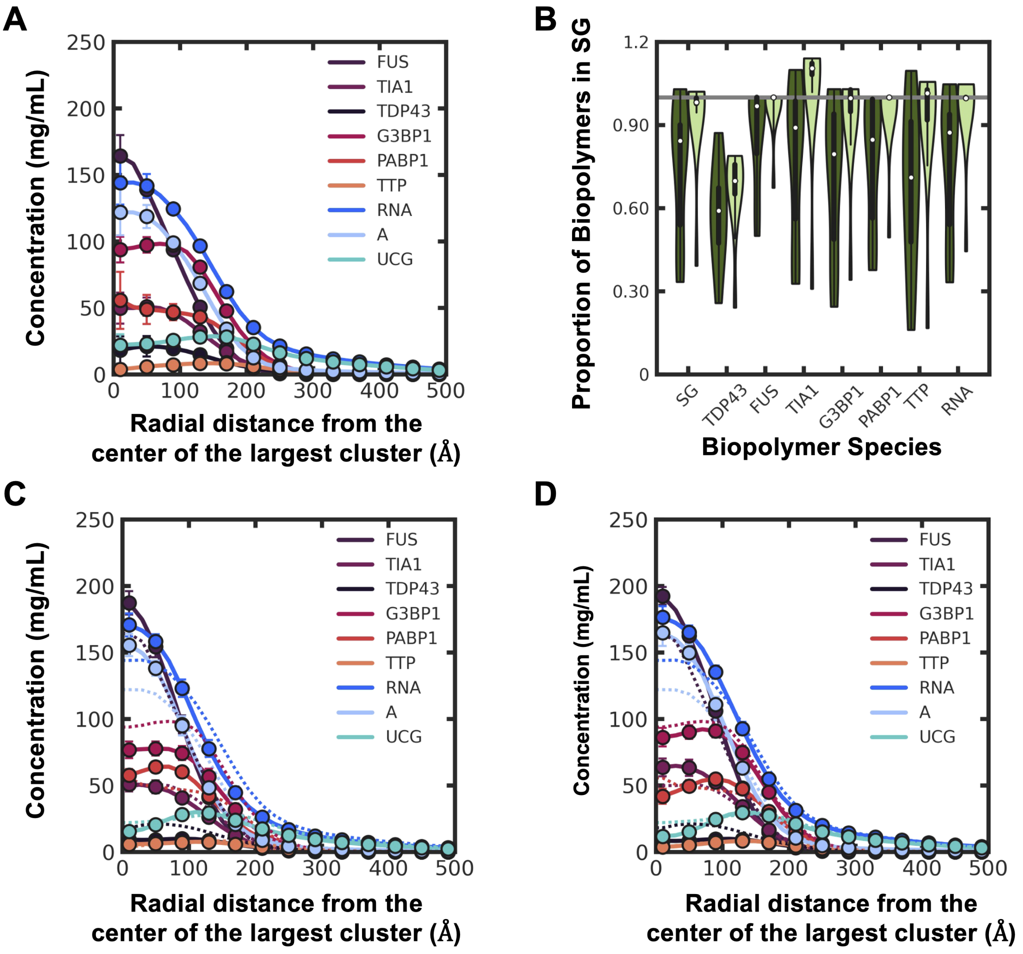


**Figure S2:** Analysis of the stress granule using radial density profiles resolved by biopolymer species and a violin plot of the change in proportion of biopolymers upon exposure to DSMs and NDSMs. **(A)** Radial mass-density profiles (RDP) resolved by biopolymer species of control stress granule. **(B)** Violin plot showing the change in proportion of biopolymers upon exposure to DSMs (dark green) and NDSMs (light green). The violin plots use the standard seaborn settings but do not extend past the most extreme data points. **(C)** Radial density profiles resolved by biopolymer species of SG systems that contain dissolving small-molecule compounds. The dashed lines are the RDPs of the control SG. **(D)** Radial density profiles resolved by biopolymer species of SG systems that contain non-dissolving small-molecule compounds. The dashed lines are the RDPs of the control SG.

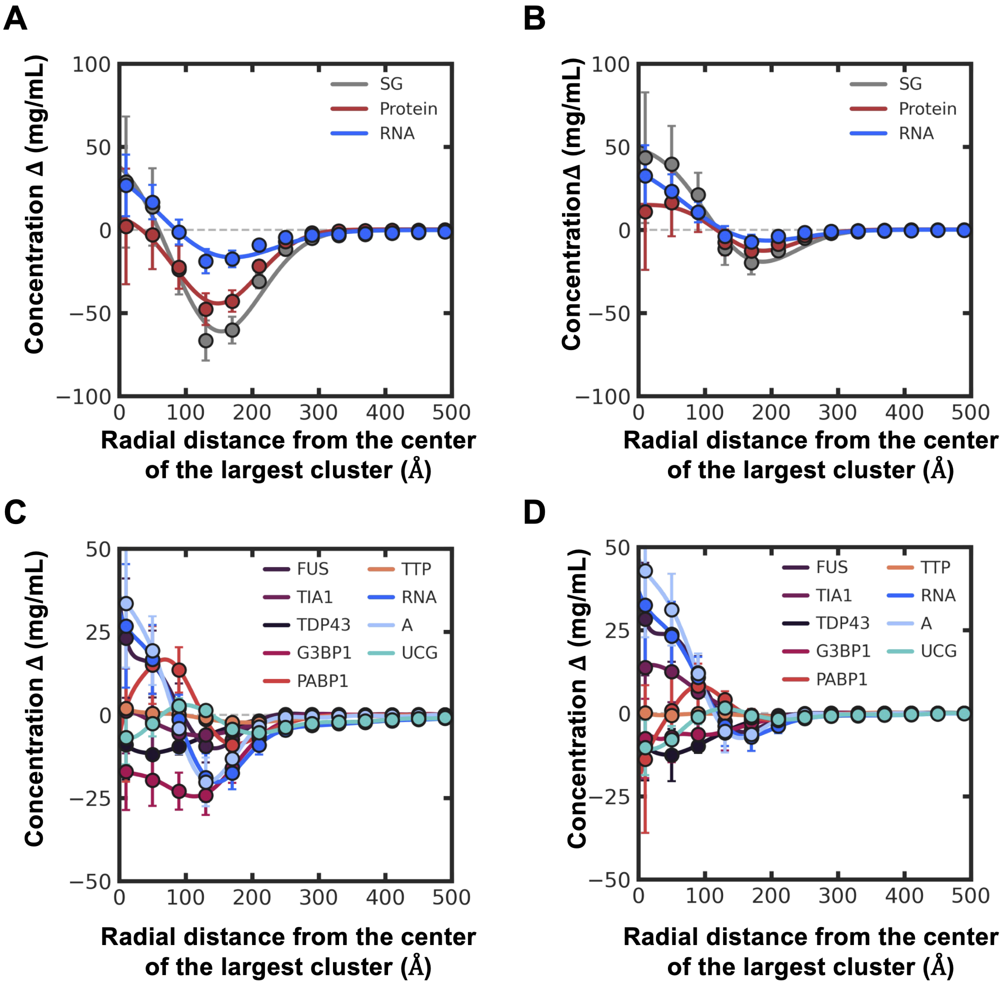


**Figure S3:** Analysis of the stress granule using the change in the radial density profiles resolved by biopolymer class and species upon exposure to DSMs and NDSMs. **(A)** The difference in the radial mass-density profiles (RDP) of the SG system resolved by biopolymer class between the control SG and the SG upon exposure to DSMs. **(B)** The difference in the radial mass-density profiles (RDP) of the SG system resolved by biopolymer class between the control SG and the SG upon exposure to NDSMs. **(C)** The difference in the radial mass-density profiles (RDP) of the SG system resolved by biopolymer species between the control SG and the SG upon exposure to DSMs. **(D)** The difference in the radial mass-density profiles (RDP) of the SG system resolved by biopolymer species between the control SG and the SG upon exposure to NDSMs.

This results in a reduced number of contacts between biopolymers and other species in systems with DSMs relative to NDSMs, as shown in Fig. S4.

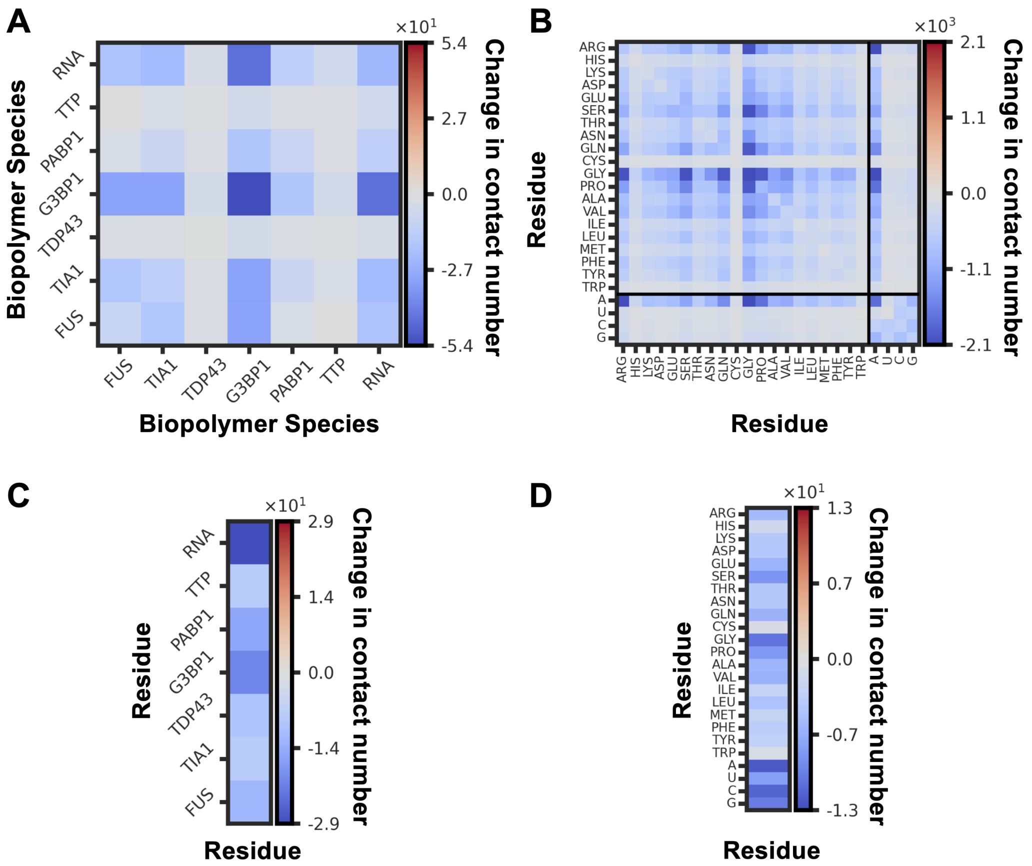


**Figure S4:** The difference in net contacts between systems that contain dissolving versus non-dissolving small-molecule compounds, reported as the change in contact number. **(A)** The difference in biopolymer–biopolymer net contacts. **(B)** The difference in amino/nucleic-acid–amino/nucleic-acid net contacts. **(C)** The difference in net contacts between biopolymers and small molecules. **(D)** The difference in net contacts between amino/nucleic acids and small molecules. Relative to non-dissolving compounds, dissolving compounds reduce biopolymer–biopolymer contacts, most strongly those involving G3BP1 and RNA.

Fig. S5 shows that there is overlap in properties between systems featuring DSM and NDSM, but the distributions possess often different means and ranges, which enables classification.

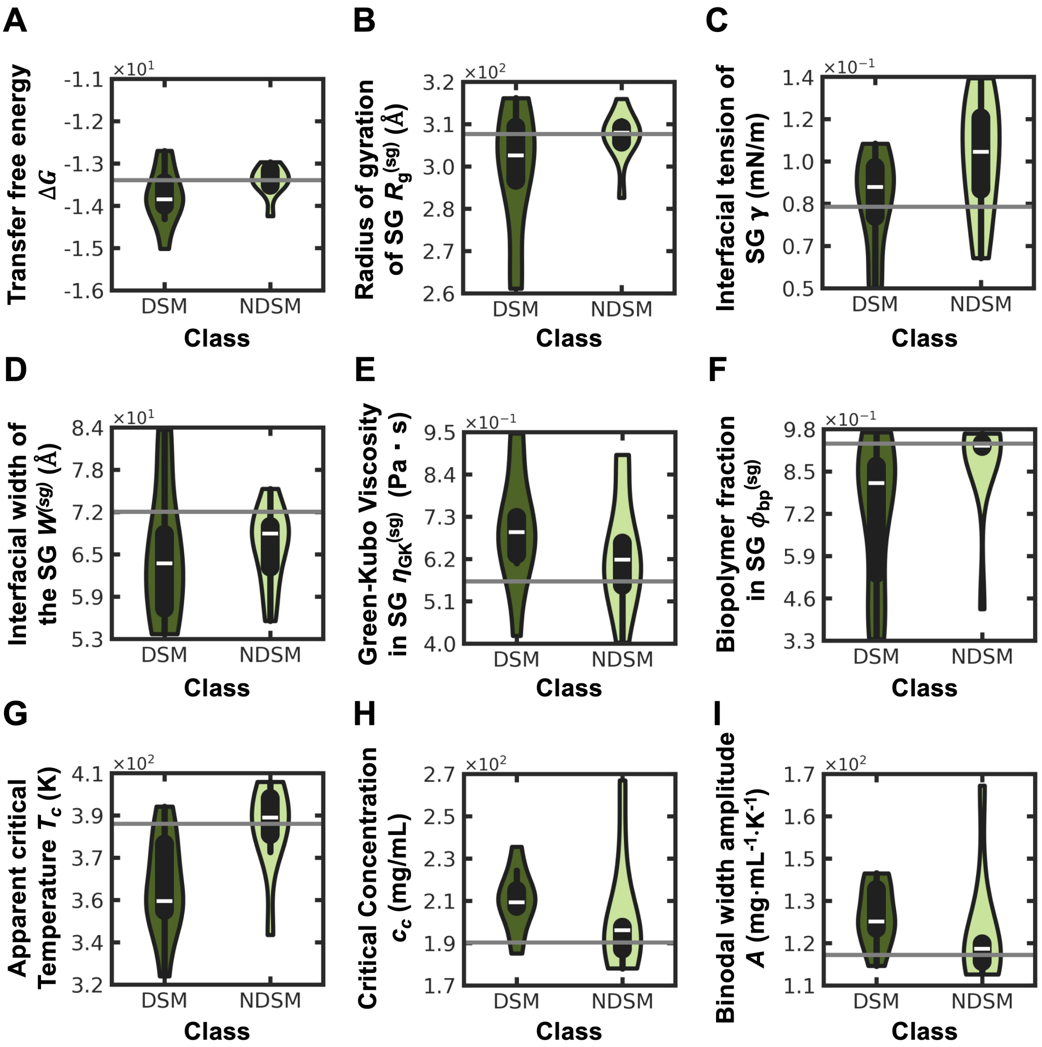


**Figure S5:** Violin-plot comparisons of the average behavior of systems containing dissolving (DSM) and non-dissolving (NDSM) small-molecule compounds for **(A)** transfer free energy $\Delta G_{\text{trans}}$, **(B)** radius of gyration $R_{\text{g}}^{(\text{sg})}$, **(C)** interfacial tension $\gamma$, **(D)** interfacial width $W^{(\text{sg})}$, **(E)** Green–Kubo viscosity $\eta_{\text{GK}}^{(\text{sg})}$, **(F)** fraction of biopolymers in the largest cluster $\phi_{\text{bp}}^{(\text{sg})}$, **(G)** apparent critical temperature $T_c$, **(H)** critical concentration $c_c$, and **(I)** binodal width amplitude $A$. The horizontal gray line marks the corresponding control SG value. The violin plots use the standard seaborn settings and do not extend past the most extreme data points.

Figs. S6 and S7 show further effects of dissolving and non-dissolving small molecules: the temporal evolution and radial structure of each system (Fig. S6) and the temperature dependence of the phase diagram and related observables (Fig. S7). In general, the dissolving and non-dissolving populations remain distinct. For the approximate binodals, the difference is more visible in the low-density regime, highlighting the instability.

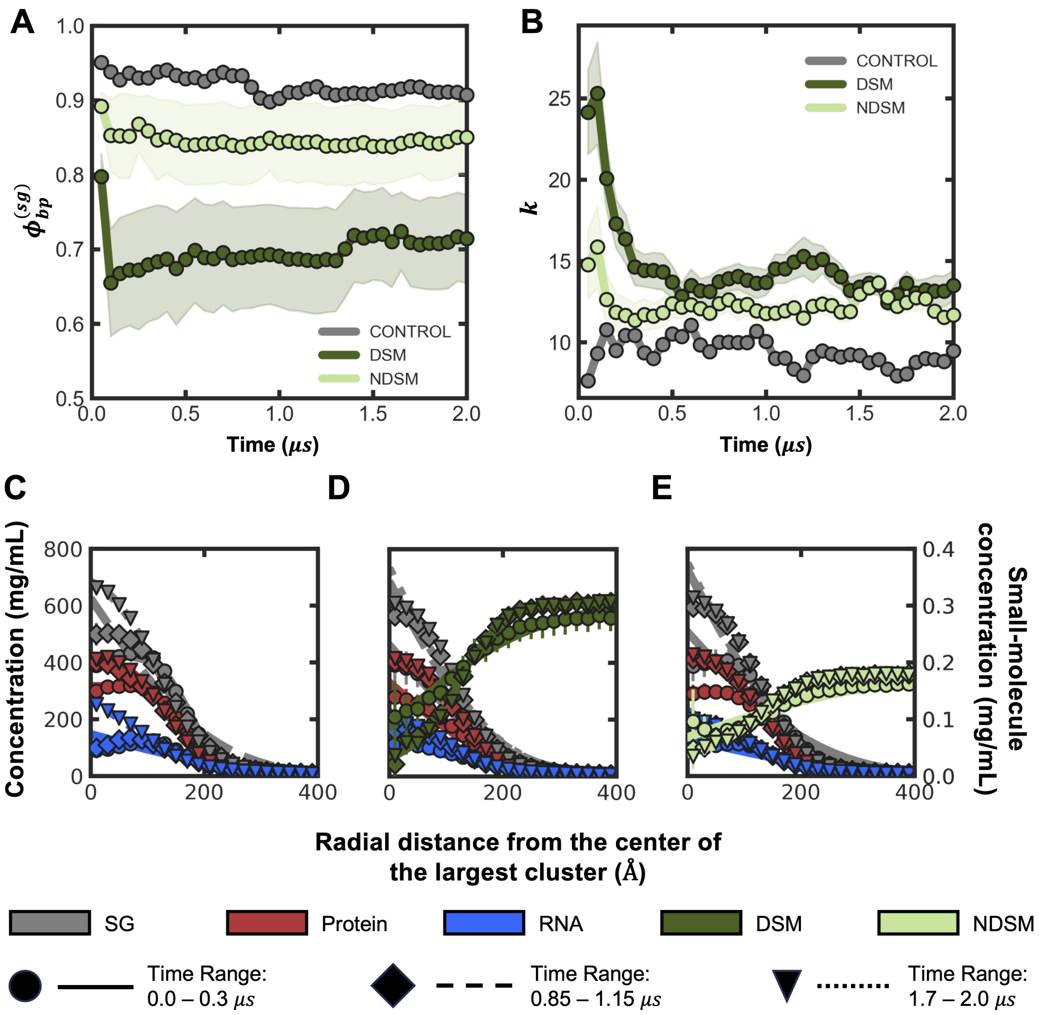


**Figure S6:** Additional comparison of Control, DSM, and NDSM systems. **(A, B)** Time evolution over the $2\ \mu$s trajectory of **(A)** the fraction of biopolymers in the largest cluster $\phi_{\text{bp}}^{(\text{sg})}$ and **(B)** the number of biopolymer clusters $k$ for the control SG, DSM, and NDSM systems; shaded bands span the ten compounds in each class. **(C–E)** Radial density profiles (concentration versus distance from the center of the largest cluster) for the **(C)** control, **(D)** DSM, and **(E)** NDSM systems, resolved into the total SG (gray), protein (red), and RNA (blue) densities, with the small-molecule concentration on the right axis (DSM dark green, NDSM light green), each shown for three time windows: early ($0.0$–$0.3\ \mu$s), middle ($0.85$–$1.15\ \mu$s), and late ($1.7$–$2.0\ \mu$s). Relative to the control and NDSMs, DSMs lower $\phi_{\text{bp}}^{(\text{sg})}$ and raise $k$.

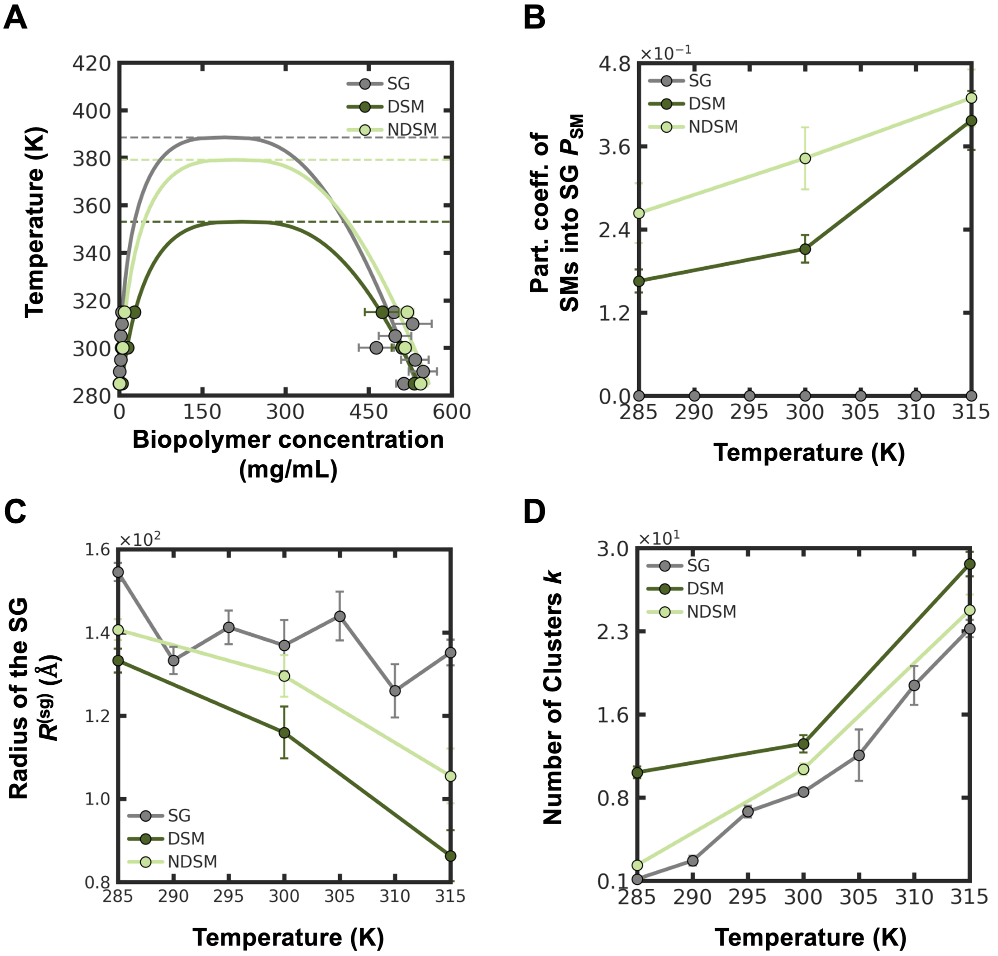


**Figure S7:** Temperature dependence across $T = 285$–$315$ K for the control SG (gray), DSM (dark green), and NDSM (light green) systems. **(A)** projected coexistence (binodal) curve in the concentration–temperature plane: markers are the dense- and dilute-phase concentrations at each temperature and the solid curves are the joint order-parameter fits, whose apexes (dashed lines) give the apparent critical temperatures ($T_c \approx 389$, $353$, and $379$ K for the control, DSM, and NDSM systems under the constrained fit). **(B)** partition coefficient of small molecules into the SG $P_{\text{SM}}$, **(C)** radius of the SG $R^{(\text{sg})}$, and **(D)** number of biopolymer clusters $k$. Relative to NDSMs, DSMs depress the apparent critical temperature and reduce the cluster size.

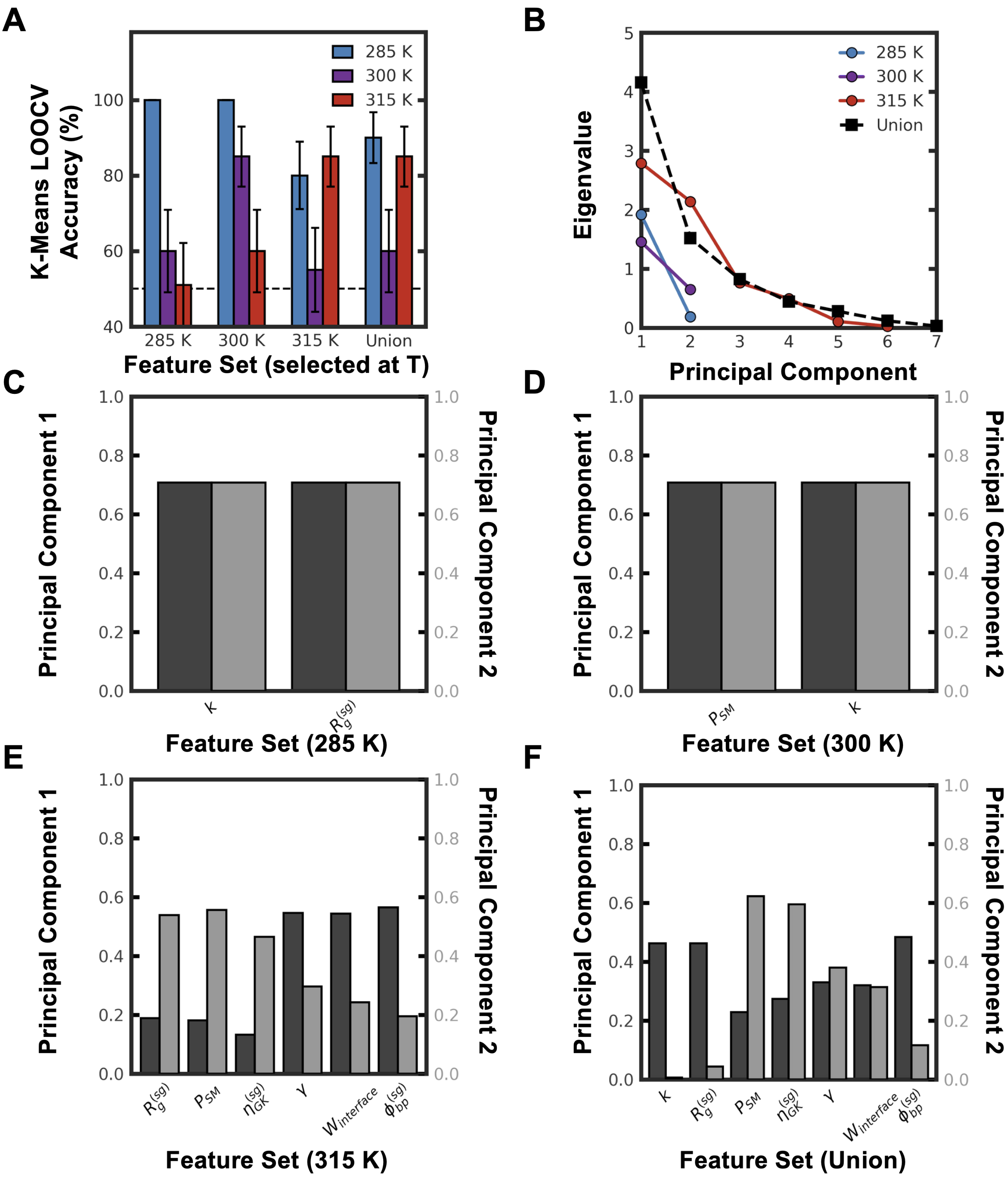


**Figure S8:** Robustness of the PCA $+\,k$-means small-molecule classifier across the three screened temperatures ($285$, $300$, and $315$ K). **(A)** Leave-one-out cross-validation accuracy when the discriminating feature set is selected at one temperature (groups) and evaluated at each temperature (colored bars); the dashed line is the $50\%$ chance level. The headline classifier of Fig. 4F is the $300$ K-selected set evaluated at $300$ K ($85\%$). **(B)** PCA eigenvalue spectra (scree) for the feature set selected at each temperature and for their union. **(C–F)** Absolute PCA loadings (left axis, first principal component, dark gray; right axis, second principal component, light gray) of the feature set selected at **(C)** $285$ K, **(D)** $300$ K (the two forward-selected features $P_{\mathrm{SM}}$ and $k$ used for Fig. 4F), **(E)** $315$ K, and **(F)** the union of all temperature-selected features.

#### S4 Temperature dependence of stress-granule structure and dissolution

To probe thermal stability, all properties were recomputed at seven temperatures spanning $285$–$315$ K (the dissolving/non-dissolving screen at the three temperatures $285$, $300$, and $315$ K). Figs. S9–S12 summarize how the control stress granule and its response to small molecules evolve across this window.

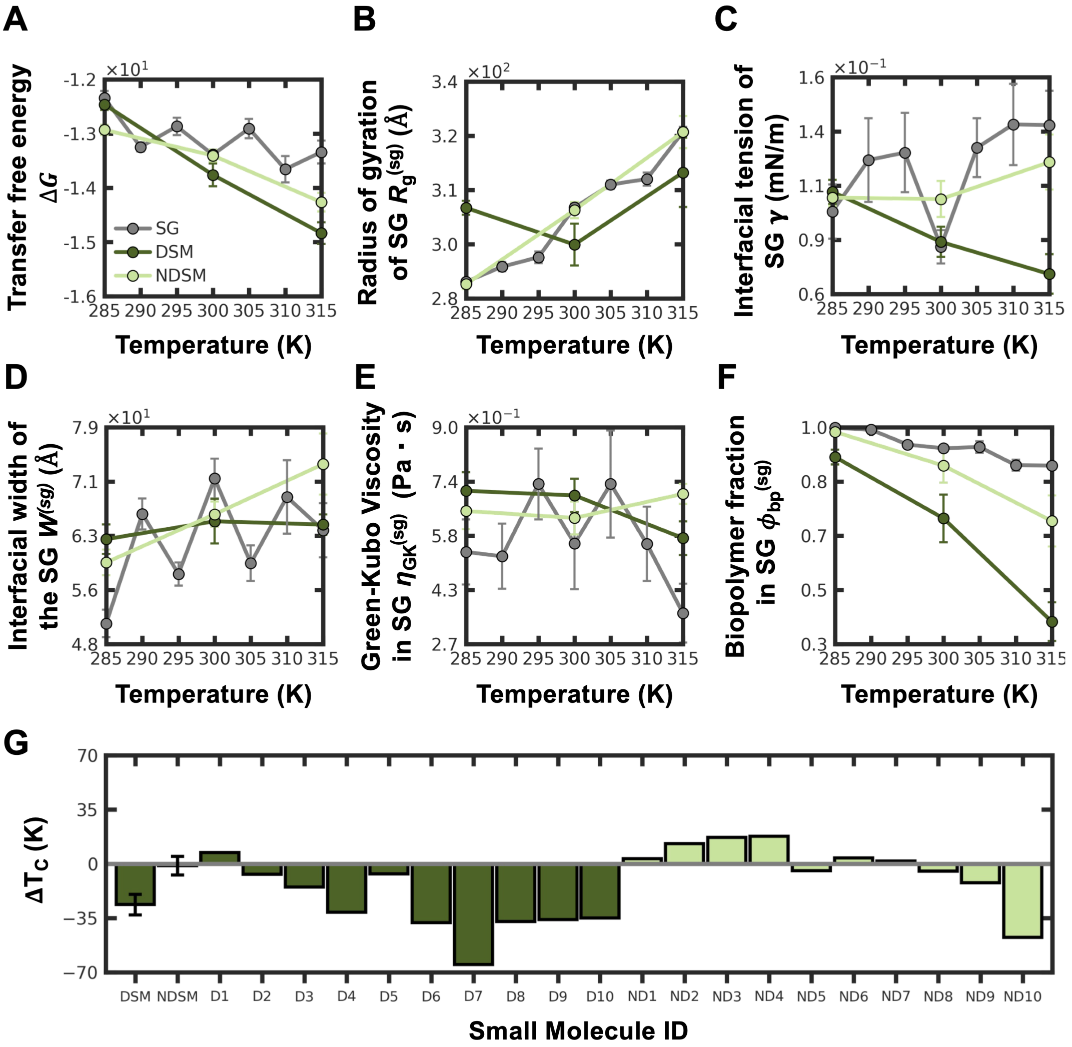


**Figure S9:** Temperature dependence of SG observables across $T = 285$–$315$ K for the control SG (gray), dissolving (DSM, dark green), and non-dissolving (NDSM, light green) small-molecule systems. **(A)** transfer free energy $\Delta G_{\text{trans}}$, **(B)** radius of gyration $R_{\text{g}}^{(\text{sg})}$, **(C)** interfacial tension $\gamma$, **(D)** interfacial width $W^{(\text{sg})}$, **(E)** Green–Kubo viscosity $\eta_{\text{GK}}^{(\text{sg})}$, and **(F)** fraction of biopolymers in the largest cluster $\phi_{\text{bp}}^{(\text{sg})}$. **(G)** apparent critical-temperature shift $\Delta T_c$ relative to the control SG for the DSM and NDSM class averages and for each individual compound (D1–D10, ND1–ND10), with DSMs in dark green and NDSMs in light green. Points in (A–F) are correlation-corrected means and error bars are $95\%$ confidence intervals. With increasing temperature the condensate loosens: the interface broadens and $\phi_{\text{bp}}^{(\text{sg})}$ falls, while the interfacial tension proxy rises modestly and $\Delta G_{\text{trans}}$ stays essentially constant. In (G) the DSM-class binodal gives a larger downward $T_c$ shift ($\approx 35$ K) than the NDSM class ($\approx 9$ K), though individual compounds overlap and D1 and ND10 disagree in sign with their experimental class.

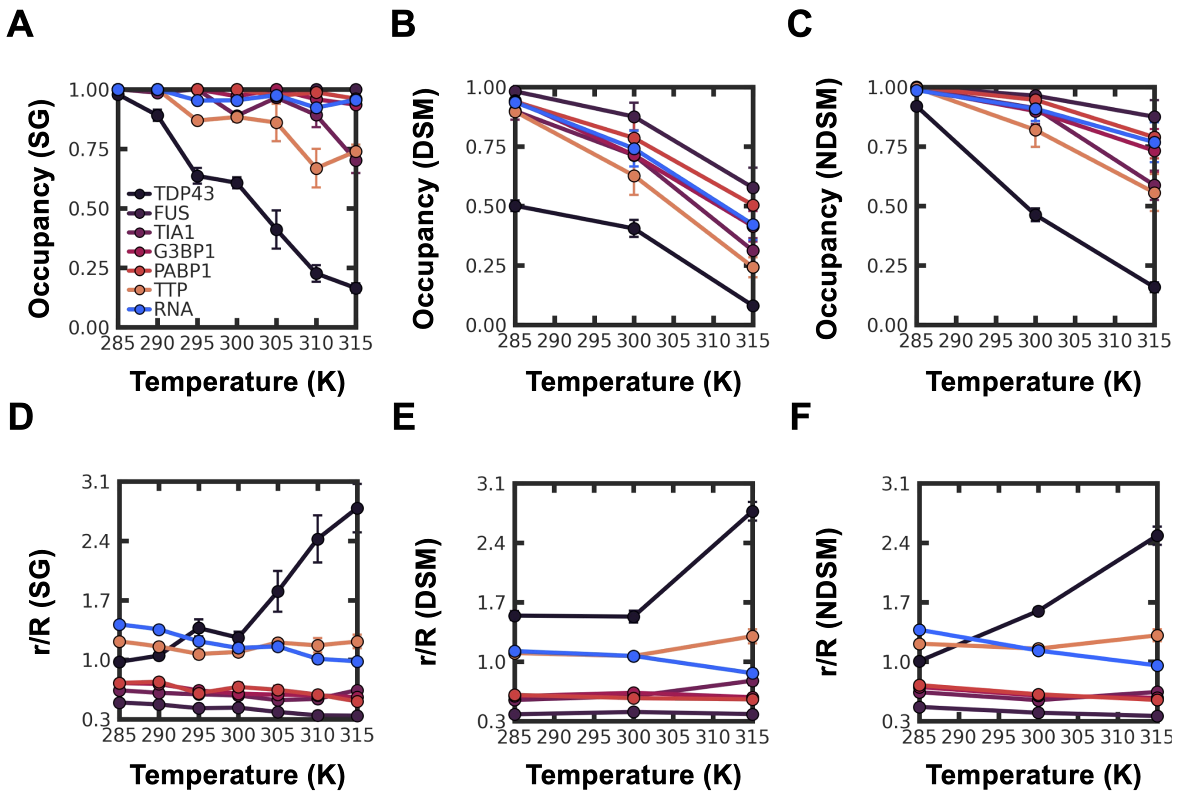


**Figure S10:** Per-species thermal response across $T = 285$–$315$ K for the control SG, dissolving (DSM), and non-dissolving (NDSM) systems. **(A–C)** condensate occupancy of each biopolymer species (TDP43, FUS, TIA1, G3BP1, PABP1, TTP, and RNA) for the **(A)** control SG, **(B)** DSM, and **(C)** NDSM systems. **(D–F)** mean radial position $r/R$ of each species, normalized by the condensate radius, for the **(D)** control SG, **(E)** DSM, and **(F)** NDSM systems. TDP43 (black) is the most thermally labile component: its occupancy collapses from near unity at $285$ K to $\approx 0.16$ at $315$ K (A) and its radial position migrates outward, from $r/R \approx 1$ at $285$ K to beyond the interface ($r/R \approx 2.7$) by $315$ K (D), whereas FUS and the remaining species stay embedded near the center ($r/R \approx 0.4$–$0.5$) at every temperature.

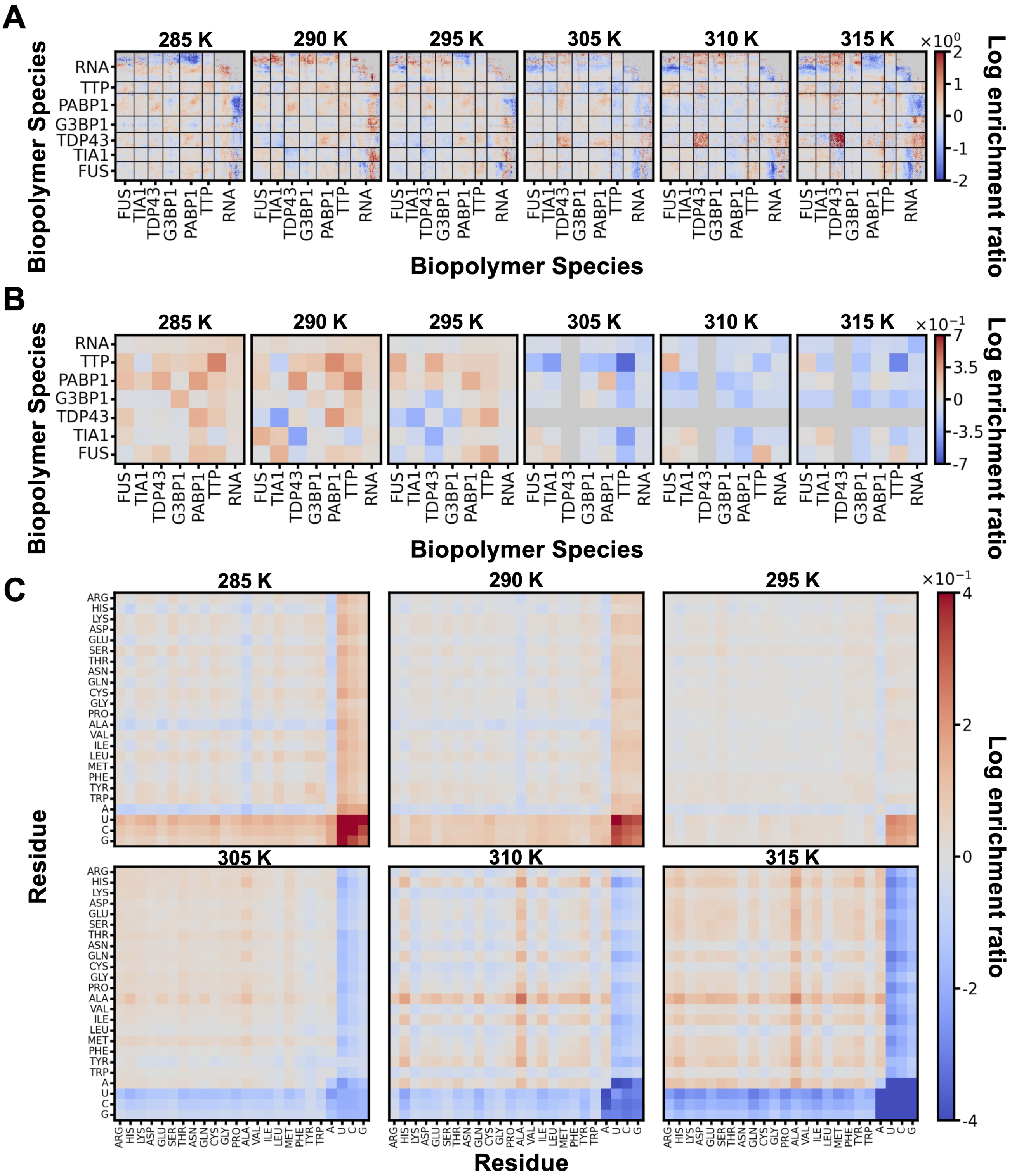


**Figure S11:** Inter-chain log contact enrichment maps of the control stress granule. Resolved by **(A)** sequence domain, **(B)** biopolymer species, and **(C)** amino/nucleic-acid type, with temperature increasing across the labeled columns from $285$ to $315$ K. Every panel is a *difference* relative to the $300$ K control: the contact enrichment at temperature $T$ minus the enrichment at $300$ K. The $300$ K reference is therefore identically zero and its column is omitted, so warm (red) cells mark pairs enriched relative to $300$ K and cool (blue) cells pairs depleted relative to $300$ K, rather than absolute enrichment at temperature $T$. The dominant interaction motifs identified at $300$ K persist across the entire window but weaken progressively as temperature increases, consistent with the falling dense-phase concentration and the broadening interface. These motifs are the G3BP1/FUS-to-RNA bridges at the species and domain level and the Arg, Gln, Tyr, and Trp contacts together with the hydrophobic Leu/Val/Ile contacts at the residue level. Contacts resolved to individual folded domains are indicative rather than quantitative (see Methods).

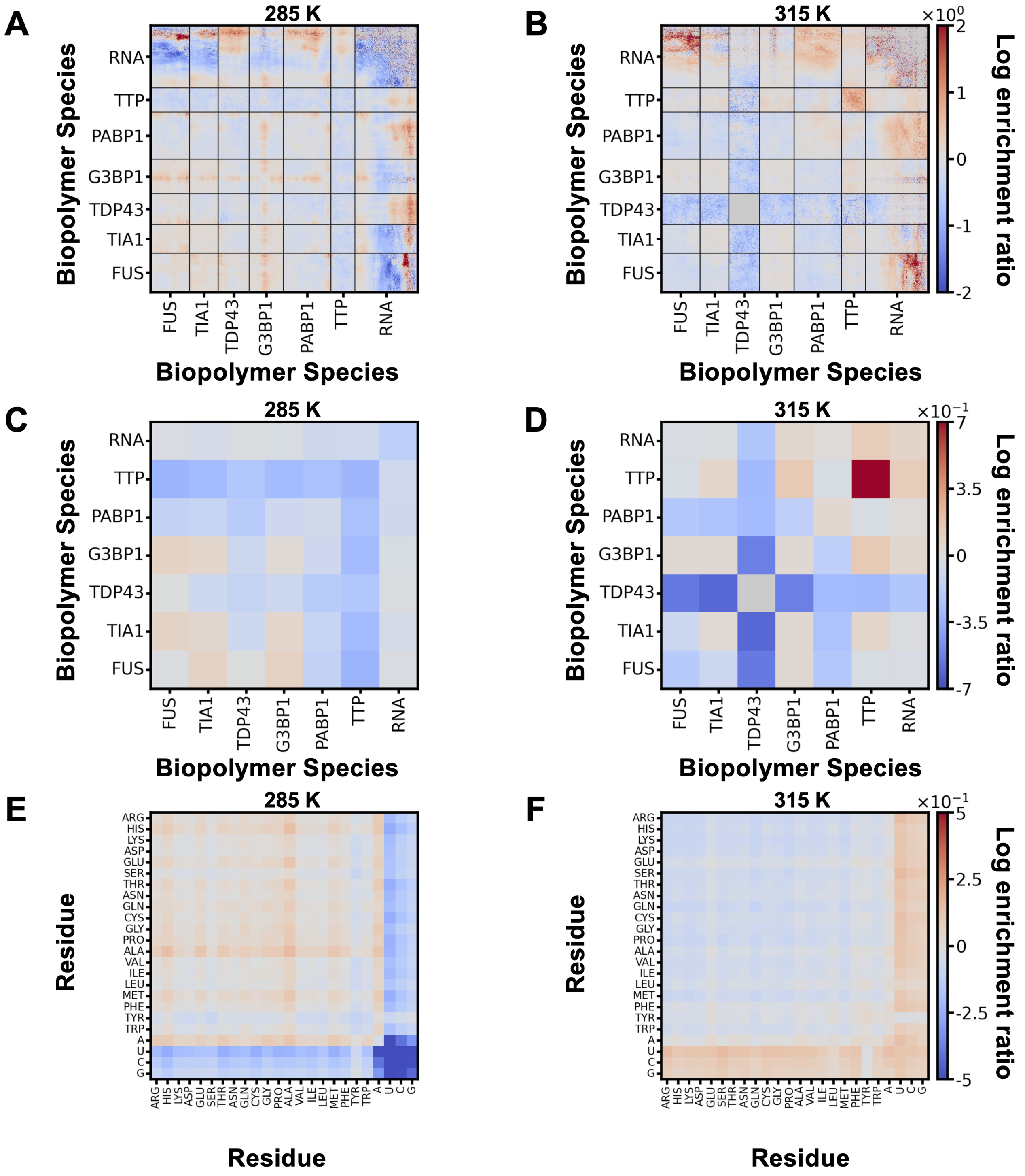


**Figure S12:** Difference (DSM-class minus NDSM-class) inter-chain contact maps, shown as the change in log contact enrichment (log enrichment ratio), resolved by sequence domain. (**A**, **B**), biopolymer species (**C**, **D**), and amino/nucleic-acid type (**E**, **F**); within each pair the left and right panels are the lowest and highest screened temperatures ($285$ and $315$ K; labeled). Each panel is additionally referenced to the $300$ K control: it shows the DSM−NDSM contrast at temperature $T$ minus the DSM−NDSM contrast at $300$ K (a double difference). The maps therefore report how the class contrast *changes* relative to $300$ K, not the raw DSM−NDSM difference at $285$ or $315$ K. The DSM−NDSM contrast in G3BP1−RNA and protein−RNA contacts grows markedly with temperature and is largest at $315$ K, reinforcing that dissolving-compound efficacy arises from a thermally assisted disruption of the RNA-scaffold contacts that hold the multiphasic architecture together. Color scales are clipped at the 99th percentile of $|\text{value}|$.

### S5 Per-species homotypic contact maps

In the pooled domain-resolved map of the main text (Figure 2C) all species share a single color scale set by the most contact-dense pair, which mutes the sparser proteins; resolved individually here, TDP43 (panel C) shows its characteristic block of homotypic contacts among the NTD, RRM, and IDR regions, confirming that this internal architecture is preserved rather than absent.

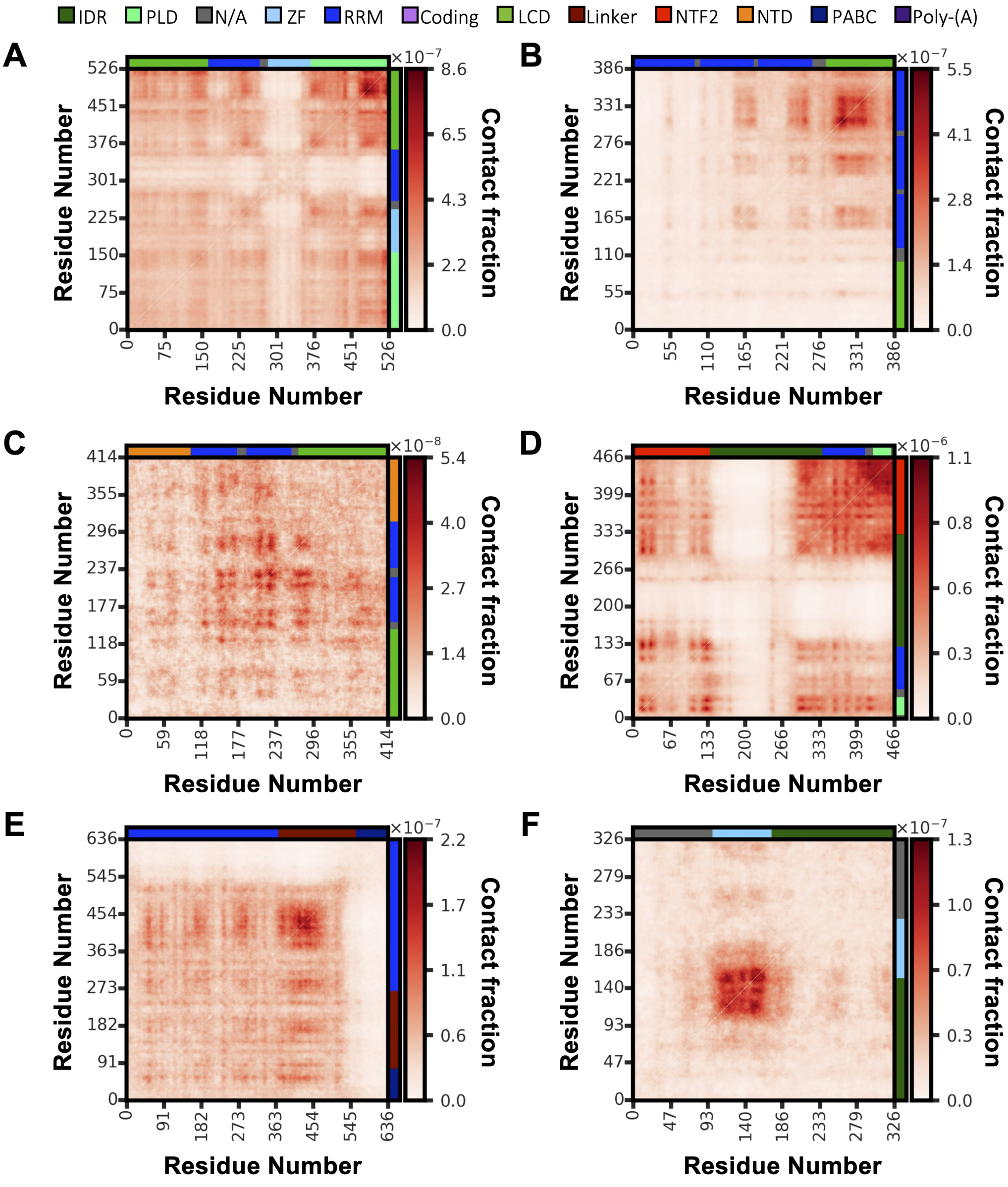


**Figure S13:** Inter-chain homotypic (self–self) contact maps for the individual protein species in the control stress granule at $300$ K, shown at single-residue resolution with the axes annotated by sequence domain (legend at top; the protein domains are IDR, PLD, LCD, Linker, NTF2, NTD, ZF, RRM, and PABC). **(A)** FUS, **(B)** TIA1, **(C)** TDP43, **(D)** G3BP1, **(E)** PABP1, and **(F)** TTP. Each panel reports the contact fraction (the share of inter-chain contacts) for residue pairs on different chains of the same species, taken from the full inter-chain contact-fraction matrix and rescaled to its own maximum, so the color-bar range differs between panels.

2. Lorentz, H.A. (1881). Ueber die anwendung des satzes vom virial in der kinetischen theorie der gase. Annalen der Physik *248*, 127–136. doi: 10.1002/andp.18812480110.

3. Berthelot, D. (1898). Sur le mélange des gaz. Comptes rendus hebdomadaires des séances de l'Académie des Sciences *126*, 1703–1855.

4. Emelianova, A., Garcia, P.L., Tan, D., and Joseph, J.A. (2025). Prediction of Small-Molecule Partitioning into Biomolecular Condensates from Simulation. JACS Au *5*, 3125–3139. URL: https://doi.org/10.1021/jacsau.5c00291. doi: 10.1021/jacsau.5c00291. Publisher: American Chemical Society.
